# A mutant form of Dmc1 that bypasses the requirement for accessory protein Mei5-Sae3 reveals independent activities of Mei5-Sae3 and Rad51 in Dmc1 filament stability

**DOI:** 10.1101/652636

**Authors:** Diedre Reitz, Jennifer Grubb, Douglas K. Bishop

## Abstract

During meiosis, homologous recombination repairs programmed DNA double-stranded breaks (DSBs). Meiotic recombination physically links the homologous chromosomes (“homologs”), creating the tension between them that is required for their segregation. The central recombinase in this process is Dmc1. Dmc1’s activity is regulated by its accessory factors Mei5-Sae3 and Rad51. We use a gain-of-function *dmc1* mutant*, dmc1-E157D*, that bypasses Mei5-Sae3 to gain insight into the role of this accessory factor and its relationship to mitotic recombinase Rad51, which also functions as a Dmc1 accessory protein during meiosis. We find that Mei5-Sae3 has a role in filament formation and stability, but not in the bias of recombination partner choice that favors homolog over sister chromatids. We also provide evidence that Mei5-Sae3 promotes Dmc1 filament formation specifically on single-stranded DNA. Analysis of meiotic recombination intermediates suggests that Mei5-Sae3 and Rad51 function independently in promoting filament stability. In spite of its ability to load onto single-stranded DNA and carry out recombination in the absence of Mei5-Sae3, recombination promoted by the Dmc1 mutant is abnormal in that it forms foci in the absence of DNA breaks, displays unusually high levels of multi-chromatid and intersister (IS) joint molecules intermediates, as well as high levels of ectopic recombination products. We use super-resolution microscopy to show that the mutant protein forms longer foci than those formed by wild-type Dmc1 (Dmc1-WT). Our data support a model in which longer filaments are more prone to engage in aberrant recombination events, suggesting that filaments lengths are normally limited by a regulatory mechanism that functions to prevent recombination-mediated genome rearrangements.

**Author Summary:** During meiosis, two rounds of division follow a single round of DNA replication to create the gametes for biparental reproduction. The first round of division requires that the homologous chromosomes become physically linked to one another to create the tension that is necessary for their segregation. This linkage is achieved through DNA recombination between the two homologous chromosomes, followed by resolution of the recombination intermediate into a crossover (CO). Central to this process is the meiosis-specific recombinase Dmc1, and its accessory factors, which provide important regulatory functions to ensure that recombination is accurate, efficient, and occurs predominantly between homologous chromosomes, and not sister chromatids. To gain insight into the regulation of Dmc1 by its accessory factors, we mutated Dmc1 such that it was no longer dependent on its accessory factor Mei5-Sae3. Our analysis reveals that Dmc1 accessory factors Mei5-Sae3 and Rad51 have independent roles in stabilizing Dmc1 filaments. Furthermore, we find that although Rad51 is required for promoting recombination between homologous chromosomes, Mei5-Sae3 is not. Lastly, we show that our Dmc1 mutant forms abnormally long filaments, and high levels of aberrant recombination intermediates and products. These findings suggest that filaments are actively maintained at short lengths to prevent deleterious genome rearrangements.

## Introduction

Homologous recombination is a high-fidelity mechanism of repair of DNA double strand breaks (DSBs), interstrand cross-links, and stalled or collapsed replication forks. In addition, during meiosis, most eukaryotes rely on CO recombination to physically link the maternal and paternal chromosomes via chiasmata, thereby making it possible for the meiosis I spindle to create the tension between homolog pairs that is required for their reductional segregation [1]. The RecA homolog Dmc1 plays the central catalytic role in meiotic recombination in budding yeast [2,3]. Following DSB formation and end resection, Dmc1 forms a helical nucleoprotein filament on the single-stranded DNA (ssDNA) tracts created by the resection machinery [4]. The nucleoprotein filament then searches the genome for a sequence of duplex DNA that is homologous to the ssDNA onto which it is loaded [5]. This region of homology can be an allelic site on one of the two homologous chromatids or on the sister chromatid. In addition, if a DSB is in a region that is repeated at more than one chromosomal locus, this can result in ectopic recombination between the two chromosomal loci [6–9]. Meiotic recombination normally favors the use of the homologous chromosome rather than the sister, consistent with the biological requirement for interhomolog (IH) COs for reductional segregation; this phenomenon is known as “IH bias” [10,11]. Once a homologous tract of double-stranded DNA (dsDNA) in found, strand exchange occurs to form a tract of hybrid DNA, pairing the ssDNA with the complementary strand of the duplex. Hybrid DNA formation displaces the opposite strand of the donor dsDNA, forming a displacement loop (D-loop) [12]. The repair process then uses the intact donor duplex DNA as a template to direct DNA repair synthesis.

Homologous recombination is highly regulated to ensure its accuracy and avoid potentially deleterious consequences of the process. Two key steps in homologous recombination, nucleoprotein filament formation and the initial invasion event, are reversible and therefore subject to this regulation [13]. Nucleoprotein filament formation, or nucleation, involves recruitment of the strand exchange protein to sites with tracts of ssDNA, nucleation of filament formation which involves displacement of the high affinity ssDNA binding protein RPA, and filament elongation which is driven by cooperative interactions between strand exchange protomers. A class of accessory proteins collectively referred to as “mediator” proteins can act to promote the displacement of RPA for filament nucleation and/or to stabilize nascent filaments, allowing them to elongate [14,15]. Mutants lacking one of these assembly proteins display defects in formation of filaments on ssDNA that can be detected by immunostaining or other cytological methods following DNA damaging treatment, or during the normal meiotic program. UvrD family helicases, including UvrD in prokaryotes and Srs2 in budding yeast, antagonize recombination at this step by disassembling ssDNA nucleoprotein filaments [16–19]. Though the strippase function of Srs2 with respect to Rad51 filaments has been well documented, Srs2 does not disassemble Dmc1 filaments, and in fact Dmc1 may inhibit Srs2 activity on ssDNA [20,21]. It is currently unknown whether there exists an ssDNA “strippase” for Dmc1.

Under normal circumstances *in vivo*, RecA family proteins form nucleoprotein filaments that are usually shorter than the resolution limit of conventional light microscopy (∼200 nanometers). This is true for RecA, and for both eukaryotic RecA homologs, Rad51 and Dmc1 [22–25]. Super-resolution microscopy imaging of Dmc1 filaments formed during meiosis indicates that Dmc1 filaments are typically about 120 nanometers long, a length that roughly corresponds to 100 nucleotides when taking into account the fact that RecA family proteins stretch the DNA ∼1.5 fold when assembled into a filament, and the length added by antibody decoration [26,27]. Furthermore, in the *exo1-D173A* mutant, in which DNA end resection is impaired during meiosis, joint molecules are formed at a level that is equivalent to wild-type, implying that short ssDNA tracts support normal meiotic recombination [28]. In addition, longer than normal Dmc1 filaments accumulate in the absence of Mnd1, a Dmc1 accessory protein that is required for Dmc1 activity after the filament formation stage [26]. Taken together, these results suggest that while RecA family proteins are competent to form long filaments, they are regulated such that they form relatively short filaments *in vivo*, though the significance of this regulation and the factors that influence filament length are not well understood.

RecA family recombinases are DNA-dependent ATPases, but their ATPase activity is not required for filament formation or for stand exchange [29–32]. Instead, ATP binding changes the conformation of the protein to a form that has high affinity for DNA, and is thus the active form [29,33]. The ADP bound form of the protein has lower affinity for DNA than the ATP-bound form, and is inactive in homology search and strand exchange. In prokaryotes, RecA ATP hydrolysis is required for filament disassembly following strand exchange, or when the protein inappropriately assembles on dsDNA [25,32]. In contrast to RecA, the eukaryotic recombinases Rad51 and Dmc1 display relatively weak intrinsic ATPase activity and rely on Rad54 family ATP-dependent dsDNA translocases to promote their dissociation [34–38]. Translocase driven dissociation is required to clear strand exchange proteins from D-loops to allow completion of recombination events [39]. Translocases also prevent accumulation of off-pathway complexes formed by filament nucleation on unbroken dsDNA [37,39–43]. The translocases may be of particular importance in eukaryotes because, unlike RecA, *in vitro* single-molecule fluorescence imaging showed that Rad51-ADP dissociation from dsDNA is inefficient and incomplete, suggesting that the activity of the translocases is required even when Rad51 is in the ADP-bound form [44]. Moreover, Rad54 was observed to have an effect on Rad51-K191R, a Rad51 mutant that is completely defective in ATP hydrolysis, implying that the ATPase activity of Rad51 is not required for it to be removed from dsDNA by Rad54 [45–47]. Finally, in the context of the nucleoprotein filament, the ATPase domain of one protomer directly contacts the N-terminal binding domain of the adjacent protomer; this observation is believed to be the structural basis for the observation that ATP-binding promotes protomer-protomer cooperativity [48,49].

We are interested in understanding how accessory proteins regulate the activity of the meiotic RecA homolog Dmc1. In *Saccharomyces cerevisiae*, Dmc1’s activity is regulated by at least five key accessory proteins including RPA, Mei5-Sae3, Hop2-Mnd1, Rad51, and the translocase Rdh54 (a.k.a. Tid1). RPA rapidly binds to tracks of ssDNA and serves to regulate interactions of Dmc1’s other accessory proteins with ssDNA [50]. *In vivo*, Mei5-Sae3 and Rad51 are required for normal Dmc1 filament formation at tracts of RPA coated ssDNA [51–53]. Hop2-Mnd1 is required for strand exchange, but not for filament nucleation or stability [54,55]. Rdh54 is a Rad54 family translocase implicated in promoting dissociation of Dmc1 from dsDNA, as discussed above.

Budding yeast Mei5-Sae3 is a homolog of *Schizosaccharomyces pombe* and mammalian Sfr1-Swi5/MEI5-SWI5, with no known homolog in plants [56]. In budding yeast, Mei5-Sae3 is Dmc1-specific, whereas in fission yeast Sfr1-Swi5 is an accessory factor to both Dmc1 and the mitotic eukaryotic RecA homolog Rad51 [57]. In mammals, MEI5-SWI5 protein is reported to function with RAD51, but there is no known interaction with DMC1, and an effort to detect DMC1 stimulatory activity *in vitro* yielded negative results [58,59]. Biochemical studies have suggested several functions for Mei5-Sae3. First, studies using fission yeast proteins have shown that Sfr1-Swi5 stimulates fission yeast Rad51, Rph51, and Dmc1 in three-stranded DNA exchange reactions, and it helps Rph51 overcome the inhibitory effect of RPA [57]. Studies using purified budding yeast Mei5-Sae3 and Dmc1 similarly concluded that Mei5-Sae3 promotes Dmc1 loading onto RPA-coated ssDNA, and that it enhances Dmc1-mediated D-loop formation when used alone, or in combination with Rad51 [3,50,60]. Haruta et al. also reported that Sfr1-Swi5 enhances Rph51’s ATPase activity; this result was subsequently clarified by work from Su et al. using purified *Mus musculus* proteins [57,59]. Su et al. showed that SWI5-MEI5 stimulates RAD51 by promoting ADP release, the step in ATP hydrolysis that is believed to be the slowest and thus rate-limiting [59,61]. Enhancement of ADP release is thought to have a stabilizing effect on Rad51 filaments by maintaining them in the ATP-bound state. In addition, later studies using a single-molecule fluorescence resonance energy transfer, concluded that mouse SWI5-MEI5 promotes RAD51 nucleation by preventing dissociation, effectively reducing the number of protomers required for a nucleation event from three to two [62]. The same study also found that fission yeast Sfr1-Swi5 prevents Rhp51 disassembly, suggesting a conserved role for this complex in stabilizing Rad51 filaments.

*In vivo*, *Saccharomyces cerevisiae* Dmc1 and Mei5-Sae3 are interdependent for focus formation, and the foci formed by the two proteins co-localize with one another, and with other DSB-dependent proteins such as Rad51 [52,53]. Moreover, Dmc1 and Mei5-Sae3 both depend on Rad51 for normal meiotic focus formation; average focus staining intensity is lower in *rad51* mutants than in wild-type [51,52]. Consistent with its requirement for Dmc1 focus formation, Mei5-Sae3 is required for Dmc1-mediated recombination *in vivo*; DSBs form normally in *mei5* or *sae3* mutants, but these intermediates are not converted to D-loops [52,53,63]. Fission yeast Rph51 differs from Dmc1 in its dependency on Sfr1-Swi5; while loss of Sfr1-Swi5 reduces recombination, recombination is only eliminated when both Sfr1-Swi5 and fission yeast Rad55-Rad57 homologs, Rph55-Rdp57, are deleted [64]. Similarly, knockdown of MEI5-SWI5 in human cells impairs RAD51 focus formation in response to ionizing radiation and also reduces recombination [58]. In contrast, deletion of mouse *Swi5* and *Sfr1* does not reduce the level of recombination when assayed with a direct-repeat reporter construct, but it does make cells more sensitive to DNA damaging agents that require homologous recombination to repair, including ionizing radiation, camptothecin, and poly(ADP-ribose) polymerase (PARP) inhibitor [65]. It is not known whether these differences in the requirement of SWI5-MEI5 by RAD51 in humans and mouse are due to differences in the cell types used or true biological differences in the human and mouse RAD51 recombinases [58].

Rad51, the RecA homolog that catalyzes homology search and strand exchange during mitotic recombination, is the second accessory protein that plays a role in forming normal Dmc1 filaments during meiosis [51]. Although Rad51 is required for normal meiotic recombination, its strand exchange activity is dispensable [3]. In fact, Rad51 strand exchange activity is inhibited during meiosis I by the meiosis-specific protein Hed1 [66,67]. In the absence of Rad51, Dmc1 foci have reduced staining intensity, suggesting that filaments are defective [51,68]. Recombination still occurs in *rad51* mutants, but it is mis-regulated such that D-loop formation occurs predominantly between sister chromatids, instead of between homologous chromosomes [69]. In addition, CO formation is reduced, only a sub-population progresses through meiotic divisions, and the spores formed are not viable [70]. In biochemical reconstitution experiments, Rad51 alone can stimulate Dmc1-mediated D-loop formation, although optimal levels of D-loop formation require both Rad51 and Mei5-Sae3 [3]. In spite of its importance as a Dmc1 accessory factor, very little is known about the molecular mechanisms involved in Rad51’s non-catalytic role in meiotic recombination. In particular, it is not known if the role of Rad51 in homolog bias is a consequence of its role in promoting Dmc1 filament formation.

One approach to studying the role of accessory proteins is to assume that the activity of the enzyme has evolved to depend on that accessory factor. In this view, beneficial regulation of an enzyme’s activity is selected for at the expense of the enzyme’s intrinsic activity. If such an evolutionary process is responsible for a particular regulatory mechanism, it should be possible to mutate the core enzyme to eliminate the “built-in” defect, rendering the mutant protein capable of catalyzing its reaction in the absence of the accessory protein. Comparison of the activities of the mutant and wild-type proteins with and without the accessory protein can then provide mechanistic insight into the processes that accessory protein normally regulates.

We applied this approach to Dmc1 in an attempt to further elucidate the mechanisms through which Mei5-Sae3 influence’s Dmc1 activity. We identified a gain-of-function Dmc1 mutant whose activity is independent of Mei5-Sae3. Characterization of this Dmc1 mutant provides new insight into the mechanism of action of Mei5-Sae3 *in vivo*, and also sheds light on the functional relationship between Mei5-Sae3 and Rad51. Furthermore, characterization of this gain-of-function version of Dmc1 reveals that it forms longer than normal filaments and displays higher than normal levels of IS, ectopic, and multi-chromatid recombination. We interpret these observations in the context of recent studies showing that a single strand exchange filament can simultaneously engage more than one dsDNA molecule.

## Results

In order to better understand the function of Mei5-Sae3 and Rad51 in Dmc1-mediated HR, we sought to identify a *DMC1* allele that would bypass the requirement for one of these accessory factors. Analysis of Dmc1-mediated recombination in the absence of an accessory factor would then allow us to identify regulatory features that depend on the accessory protein by comparison to the wild-type process. To this end, we constructed two *dmc1* mutants based on two previously characterized gain-of-function mutations in Dmc1 homologs, RecA-E96D and Rad51-I345T [32,71,72]. Sequence alignments indicated that the amino acid residues altered in these mutants are conserved allowing us to construct corresponding mutant forms of Dmc1; for RecA-E96D the corresponding mutant is Dmc1-E157D and for Rad51-I134T the corresponding mutant is Dmc1-I282T.

To assess whether either of these Dmc1 mutants would bypass Mei5-Sae3 and/or Rad51, we constructed diploid yeast lacking either Mei5 or Rad51 with the corresponding Dmc1 mutation, and assessed sporulation efficiency and spore viability alongside *DMC1^+^ mei5* and *DMC1^+^ rad51* controls. In a *mei5* strain, tetrads are formed very inefficiently, whereas in a *rad51* mutant, tetrads are formed, but almost all spores within them are dead [52,53,70]. We found that *dmc1-E157D* bypasses Mei5-Sae3 with respect to sporulation and spore viability (Table 1). The spore viabilities of *dmc1-E157D*, *dmc1-E157D mei5*, and *dmc1-E157D sae3* are nearly identical to one another (57.6%, 50.3%, and 57.0% respectively), suggesting that Dmc1-E157D function is largely independent of Mei5-Sae3. In contrast, *dmc1-E157D* does not bypass the requirement for *rad51* with respect to spore viability (0.0% in *rad51* versus 0.74% in *dmc1-E157D rad51*).

**Table 1.**
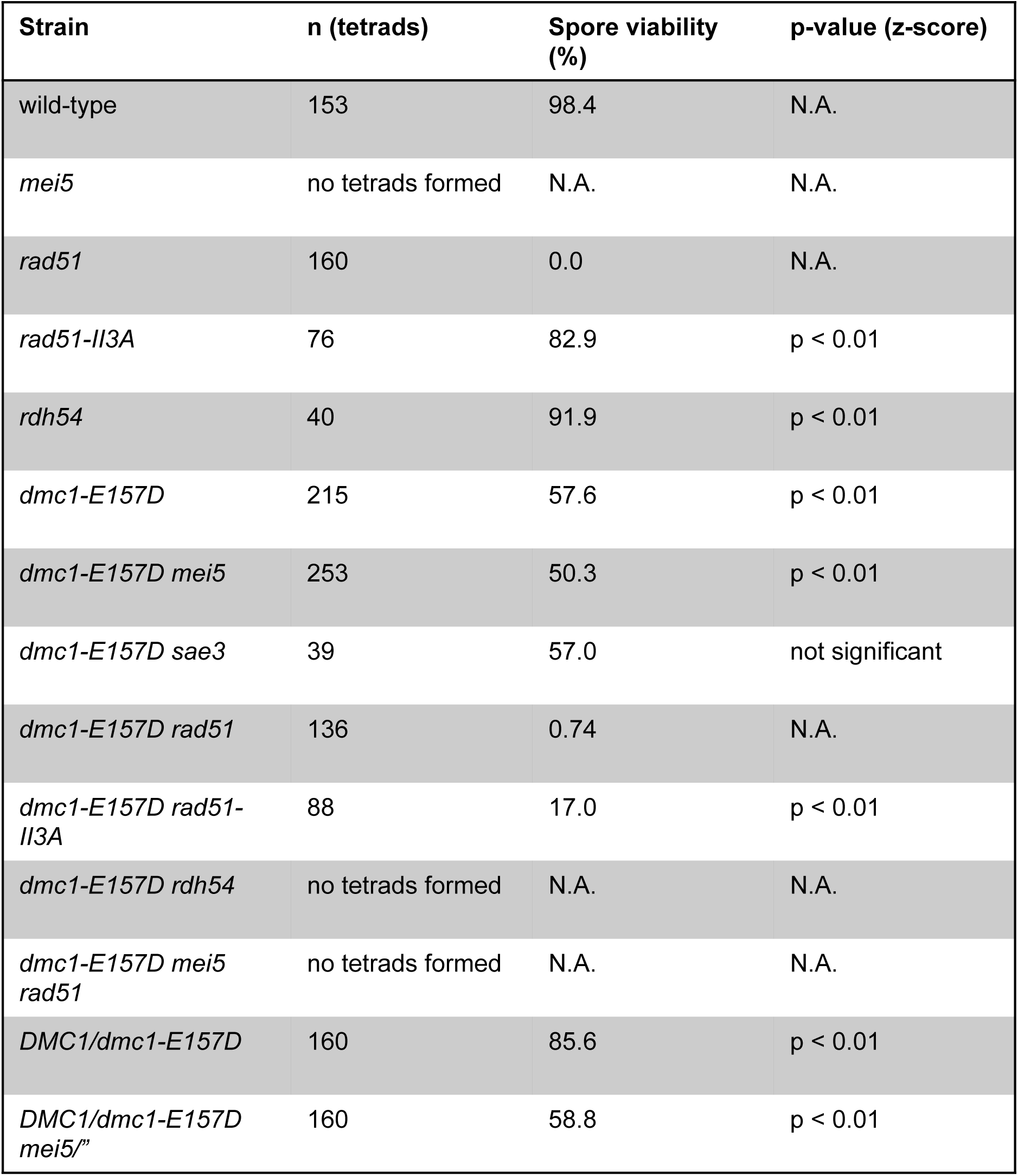
Spore viabilities for strains in study. p-values are reported for z-scores. Comparison for single mutants is to wild-type. Comparison for double mutants is to each of the single mutants; the largest p-value is reported. Comparison for heterozygotes is to homozygotes. N.A. = not applicable; for samples that do not meet the success/failure condition for z-scores and wild-type to itself. Strains used in experiments in the order in which they appear in table, top to bottom: DKB3698, DKB6320, DKB3710, DKB3689, DKB2526, DKB6342, DKB6299, DKB6300, DKB6539, DKB6540, DKB6393, DKB6400, DKB6583, DKB6412, DKB6413, DKB6525, DKB6619, DKB6406, DKB6407.

Spore viability data from the *dmc1-E157D/DMC1* heterozygote and *dmc1-E157D/DMC1 mei5/”* heterozygote strains suggests that Dmc1-E157D is co-dominant with wild-type Dmc1 in the presence of Mei5 (85.6%), but fully dominant to wild-type Dmc1 in the absence of Mei5 (58.8% in *dmc1-E157D/DMC1 mei5/”* versus 50.3% in *dmc1-E157D mei5*). In contrast to *dmc1-E157D*, we did not detect phenotypic suppression in *dmc1-I282T* mutants, either with respect to prophase arrest in a *mei5* mutant background, or with respect to spore viability in a *rad51* background. Importantly, the Dmc1-E157D mutation does not result in increased expression or stability of the protein as assayed by Western blotting of meiotic yeast whole cell extracts, thus ruling out a trivial explanation for Dmc1-E157D’s bypass of the *mei5* and *sae3* mutations (Supplemental Figure 1).

### Dmc1-E157D forms meiotic immunostaining foci in the absence of Mei5 and Rad51

We next performed immunofluorescence staining of spread meiotic nuclei to examine Dmc1 focus formation in the *dmc1-E157D* and *dmc1-E157D mei5* strains. As shown previously, meiotic Dmc1-WT focus formation is severely defective in *mei5* mutant cells, but *dmc1-E157D* forms bright Dmc1 foci in the *mei5* mutant background (Figures 1a,b) [52,53]. Notably, Dmc1 foci accumulate to higher levels and persist for longer in *dmc1-E157D* and *dmc1-E157D mei5* when compared to wild-type.

One model suggests that Mei5-Sae3 and Rad51 cooperate to promote Dmc1 filament formation. Because *dmc1-E157D* bypasses *mei5*, we reasoned that if this model is correct, *dmc1-E157D* might also bypass the defect seen for formation of brightly-staining Dmc1 foci in *rad51* cells, even it does not suppress the spore viability defect observed in these cells. To test this, we constructed *dmc1-E157D rad51* and *dmc1-E157D mei5 rad51* strains, and looked for Dmc1 focus formation in spread meiotic nuclei. In contrast to a *rad51* single mutant, in which Dmc1-WT staining intensity is reduced, the Dmc1 foci observed in *dmc1-E157D rad51* and *dmc1-E157D mei5 rad51* nuclei were brighter and more numerous than those in wild-type (Figures 1c,d) [51,68]. We conclude that *dmc1-E157D* bypasses the role of Rad51 with respect to forming brightly staining Dmc1 foci.

### *dmc1-E157D* forms immunostaining foci in the absence of DSBs

Because the Dmc1-E157D mutant is modeled after RecA-E96D, which has been shown to form foci on undamaged DNA, we wanted to ask whether the same was true of the corresponding Dmc1 mutant [25]. To determine whether any of the foci that we observed in the *dmc1-E157D* background resulted from binding to chromosomes independent of DSBs, we introduced the *spo11* mutation into our *dmc1-E157D* strains to block DSB formation. Spo11 is the catalytic subunit of a meiosis-specific complex that induces DSBs at the outset of meiosis [73]. Immunostaining of spread meiotic nuclei for Dmc1 and RPA revealed that in contrast to the *spo11* single mutant, which typically forms few if any Dmc1 foci, nearly all *spo11 dmc1-E157D* nuclei contained numerous Dmc1 foci (Figures 1e,f) [41]. RPA serves as a marker for DSB-associated tracts of ssDNA in mid-to-late prophase I. RPA foci are detected early in prophase in *spo11* mutants owing to the role of RPA in pre-meiotic DNA replication, but then disappear 4 hours after induction of meiosis [74]. We found that at 4 hours, the majority of nuclei lacking RPA foci contained Dmc1 foci in *spo11 dmc1-E157D* and *spo11 dmc1-E157D mei5* (100% and 96% of nuclei lacking RPA had Dmc1 foci, respectively) (Figure 1e). We conclude that Dmc1-E157D forms DSB-independent foci, suggesting that a substantial fraction of the foci observed in *SPO11^+^ dmc1-E157D* cells represent off-pathway structures formed by binding unbroken chromosomal loci.

**Figure 1.**
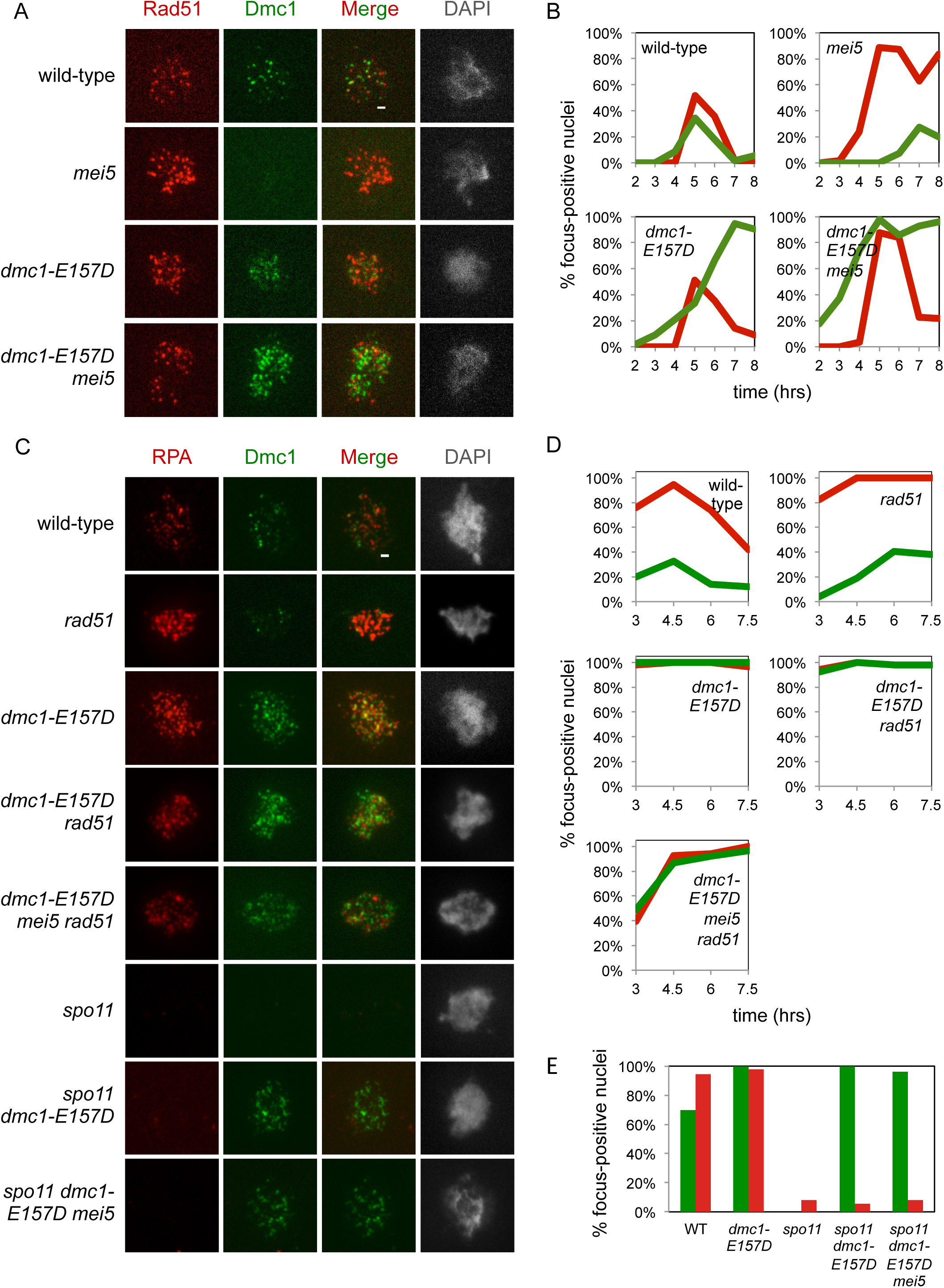
*dmc1-E157D* bypasses *mei5*, *rad51* with respect to focus formation. (a, c) Representative widefield microscopy imaging of spread meiotic nuclei are shown for each strain. Scale bars represent 1 µm. (b, d) Quantitation. Nuclei were scored as focus positive if they contained three or more foci of a given type. Dmc1 (green), Rad51 or RPA (red). (e) Quantitation of *spo11* strains and controls at 4 hours. Strains used in experiments in the order in which they appear in figure, top to bottom: DKB3698, DKB6320, DKB6342, DKB6300, DKB3710, DKB6393, DKB6412.

### *dmc1-E157D* bypasses Mei5, but not Rad51, with respect to meiotic CO formation

To examine whether Dmc1-E157D is competent to carry out recombination in the absence of Mei5, we performed 1D gel electrophoresis, followed by Southern blotting, to detect DNA double strand breaks and formation of CO recombination products at the well-characterized recombination hotspot *HIS4::LEU2* [75]. 1D gel electrophoresis at *HIS4::LEU2* can be used to detect DSB intermediates and IH CO products. In addition, the same gels detect products that result from ectopic recombination between the *HIS4::LEU2* locus and the native *LEU2* locus, which are separated by ∼23 kilobases on chromosome III [9,76]. As shown previously, DSBs accumulate and CO formation is very limited in *DMC1^+^ mei5* (Figures 2a,b) [52,53]. In contrast, although *dmc1-E157D mei5* cells accumulate DSBs, these intermediates are resolved by 24 hours, at which point CO formation is equivalent to wild-type (Figures 2a,b). In addition, whereas only 8.7% of *DMC1^+^ mei5* cells progress through a meiotic division, 50.0% of *dmc1-E157D mei5* cells progress, a level nearly equivalent to *dmc1-E157D* (58.4%) (Figure 2b). This shows that Dmc1-E157D bypasses the normal requirement for Mei5 during meiotic recombination. Interestingly, ectopic recombination is elevated ∼3.5-fold in *dmc1-E157D* and *dmc1-E157D mei5* relative to wild-type.

**Figure 2.**
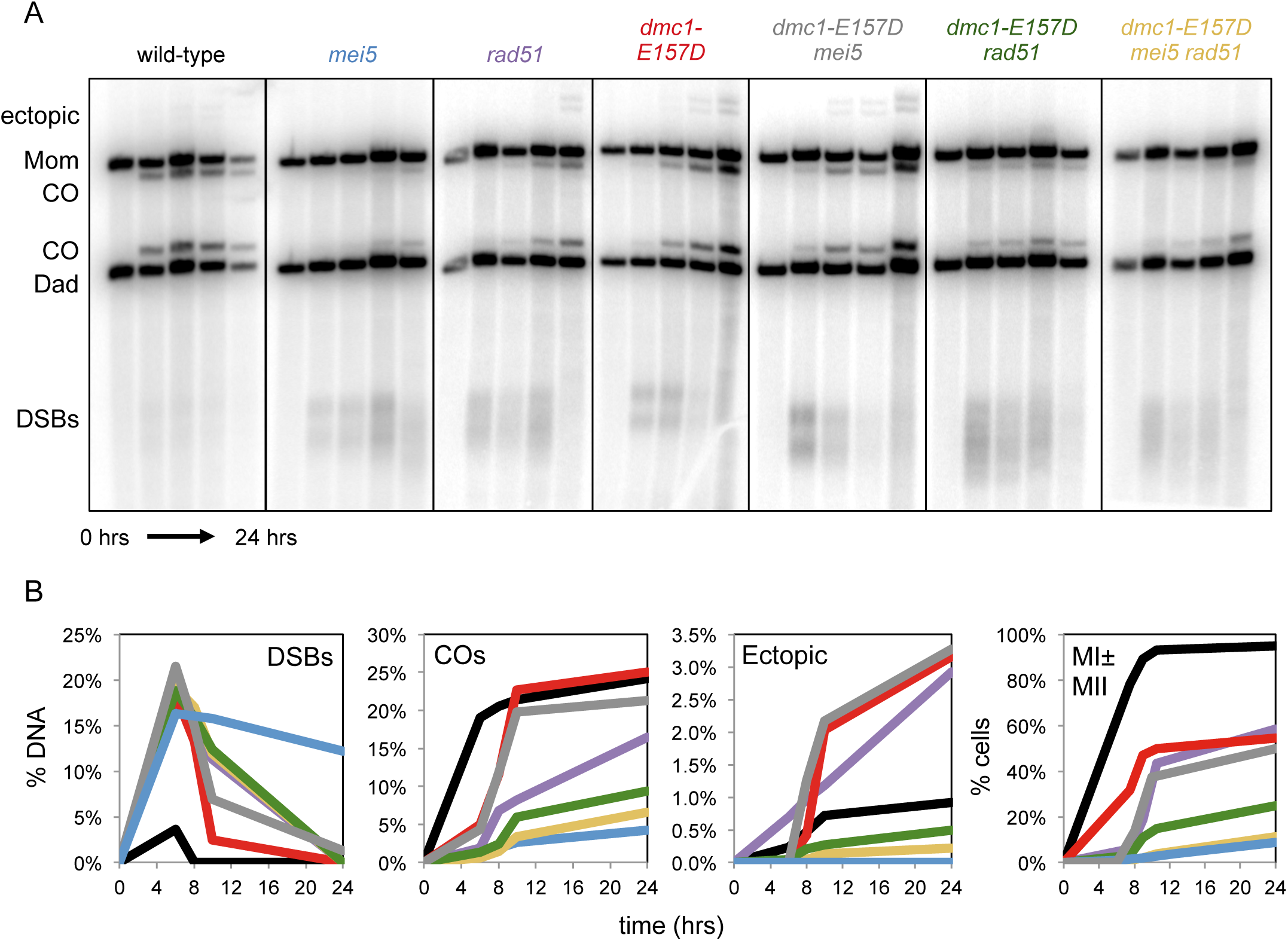
*dmc1-E157D* bypasses *mei5* but not *rad51* with respect to CO formation. (a) 1D XhoI gels at the *HIS4::LEU2* hotspot from meiotic time course experiments (b) Quantitation of 1D gels shown in (a); black – wild-type, light blue – *mei5*, purple – *rad51*, red – *dmc1-E157D*, gray – *dmc1-E157D mei5,* green – *dmc1-E157D rad51*, yellow – *dmc1-E157D mei5 rad51* and meiotic progression data for each strain. For each time point, ≥50 cells were scored. Strains used in experiments in the order in which they appear in figure, right to left: DKB3698, DKB6320, DKB3710, DKB6342, DKB6300, DKB6393, DKB6412.

Because Dmc1-E157D also bypasses Rad51 with respect to forming brightly staining foci, we wanted to ask whether it similarly bypasses Rad51 for CO formation and DSB resolution at *HIS4::LEU2*. Previous studies of *rad51* mutants showed that DSBs accumulate and undergo more extensive resection than wild-type [70]. In addition, the final level of COs that form in *rad51* was reported to be 5-fold lower than wild-type, and ectopic recombination is ∼1.6-fold higher at 10 hours in sporulation medium [70,77]. We confirmed these phenotypes for the *rad51* single mutant (Figures 2a,b). Consistent with the failure of *dmc1-E157D* to rescue the low spore viability phenotype of *rad51*, we found that *dmc1-E157D rad51* accumulates hyper-resected DSBs (Figure 2a). Surprisingly, *dmc1-E157D rad51* makes fewer COs than *rad51*, implying that the *dmc1-E157D rad51* double mutant is more defective than either the *dmc1-E157D* single mutant or the *rad51* single mutant. Meiotic progression data similarly indicates that *dmc1-E157D* is more defective than both *dmc1-E157D* and *rad51*; only 24.8% of *dmc1-E157D rad51* cells execute at least one meiotic division, compared to 58.4% and 54.5% of *dmc1-E157D* cells and *rad51* cells respectively (Figure 2b). Additionally, very little ectopic recombination is detected in *dmc1-E157D rad51*, possibly reflecting the fact that there is less recombination overall, or a change in the pattern of formation of joint molecules (JMs) or how they are resolved. Overall our results indicate that *dmc1-E157D* does not bypass *rad51* with respect to resolution of meiotic DSBs.

The *dmc1-E157D mei5 rad51* triple mutant was similar to the *dmc1-E157D rad51* double, with the triple displaying slightly more pronounced defects final CO levels (Figure 2a,b). We also found that the efficiency of the first meiotic division is somewhat reduced in the *dmc1-E157D mei5 rad51* mutant (11.3%) compared to the *dmc1-E157D rad51* (24.8%) mutant (Figure 2b). These results indicate that recombination in *dmc1-E157D* displays a strong dependence on Rad51, but no dependence on Mei5, unless Rad51 is absent.

Introduction of the *spo11* mutation into the *dmc1-E157D* and *dmc1-E157D mei5* backgrounds rescued the meiotic progression defects observed for *dmc1-E157D* and *dmc1-E157D mei5* (Supplemental Figure 2). Thus the meiotic progression defects observed in the *dmc1-E157D* background are DSB-dependent. This finding suggests that although there are numerous DSB-independent Dmc1 foci in these strains, these Dmc1-dsDNA complexes do not dramatically interfere with chromosome segregation.

### Dmc1-mediated meiotic recombination is independent of Mei5-Sae3 in *dmc1-E157D*

We next sought to further characterize Dmc1-E157D-mediated recombination in the absence of Mei5 by 2D gel electrophoresis and Southern blotting. Using this method, an array of JM recombination intermediates can be detected at the *HIS4::LEU2* locus, including single-end invasions (SEIs), IS-double Holliday junctions (IS-dHJs), IH-double Holliday junctions (IH-dHJs), and multi-chromatid joint molecules (mcJMs) [78]. Representative 2D gel images are shown for each strain in Figure 3a. As expected, JM formation is severely compromised in *mei5* (Figures 3a,b). In contrast, in *dmc1-E157D mei5*, JM formation is efficient, with IH-dHJ levels equivalent to those in wild-type. IS-dHJs, however, are increased ∼3-fold, reducing the IH-dHJ/IS-dHJ ratio from 5.0 in wild-type to ∼1.5 in *dmc1-E157D* (Figure 3b). *dmc1-E157D mei5* phenocopies *dmc1-E157D*, which also has increased IS-dHJs and a reduced IH/IS ratio of ∼1.6. SEIs are observed at the same levels in *dmc1-E157D* and *dmc1-E157D mei5* mutants as in wild-type (Figure 3b). Like IS-dHJs, mcJMs are increased relative to wild-type in both *dmc1-E157D* and *dmc1-E157D mei5* (3.0-fold and 2.7-fold respectively). The similar array of JMs observed in *dmc1-E157D* and *dmc1-E157D mei5* cells further indicates that Dmc1-E157D-mediated recombination occurs independently of Mei5-Sae3. Although a decrease in the IH/IS ratio can be interpreted as a defect in the mechanism of IH bias, this case is unusual in that the decreased ratio results from increased IS-dHJs, with no reciprocal decrease in IH-dHJS. The fact that the level of IH-dHJs in *dmc1-E157D mei5* cells is equivalent to that in wild-type suggests that the mechanism of homolog bias is intact in this mutant, and reveals that Mei5-Sae3 is not required for IH bias. The data also suggest that the *dmc1-E157D* mutant is a hyper-recombinant mutant, displaying higher than normal levels of IS-dHJs and mcJMs, as well as increased ectopic COs.

**Figure 3.**
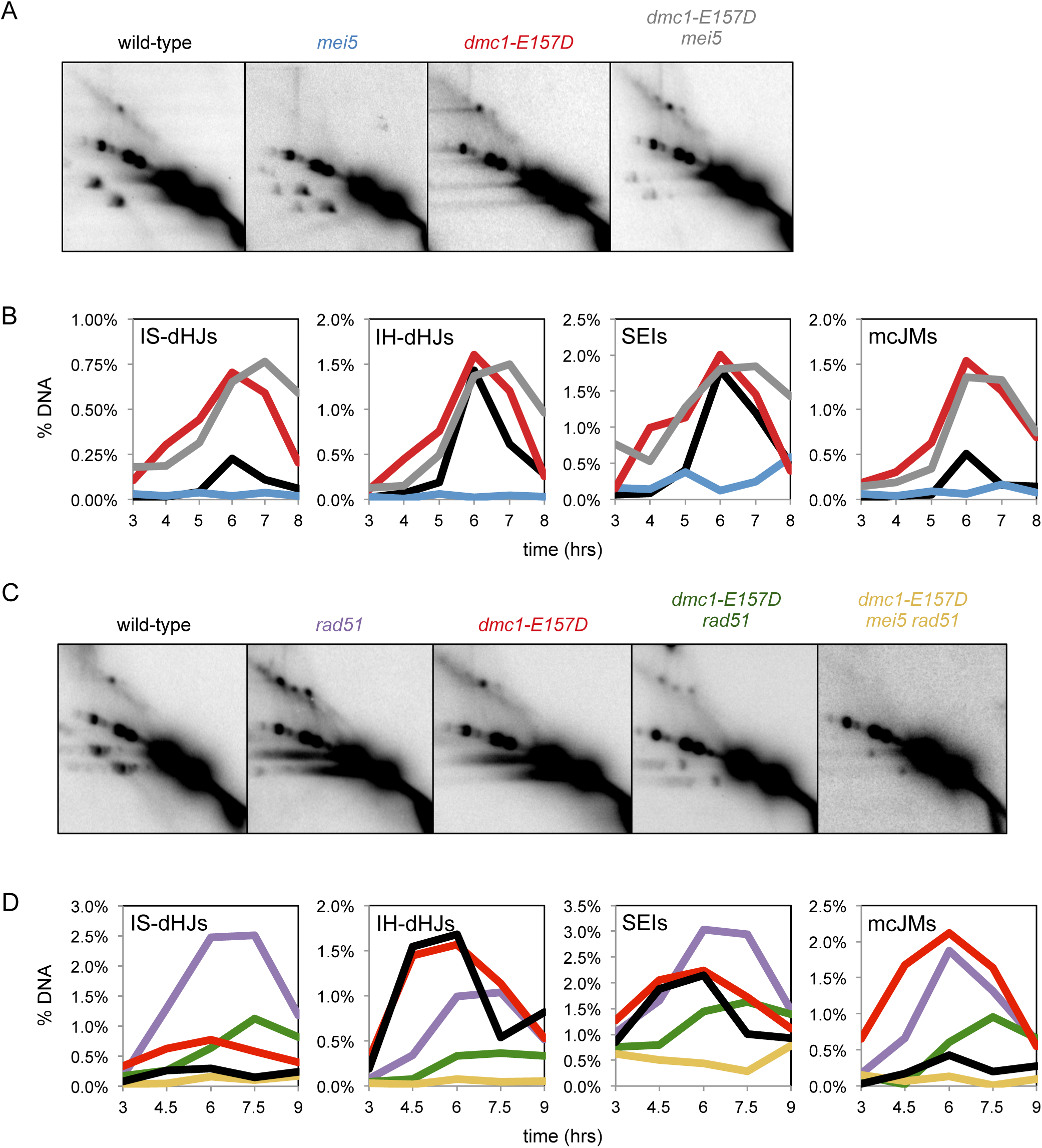
Recombination in *dmc1-E157D* is abnormal and dependent on Rad51, with little affect of Mei5-Sae3. (a) 2D gels at the *HIS4::LEU2* hotspot from meiotic time course experiments. Representative images were chosen according to the time at which total JMs peaks for each sample (wild-type, 6 hours; *mei5*, 8 hours; *dmc1-E157D*, 8 hours; *dmc1-E157D mei5,* 7 hours) (b) 2D gel quantitation; black – wild-type, light blue – *mei5Δ*, red – *dmc1-E157D*, gray – *dmc1-E157D mei5*. (c) 2D gels. Representative images were chosen according to the time at which total JMs peaks for each sample (*rad51,* 6 hours; *dmc1-E157D,* 6 hours; *dmc1-E157D rad51*, 7.5 hours; *dmc1-E157D mei5 rad51*, 6 hours) (d) 2D gel quantitation; black – wild-type, light purple – *rad51*, red – *dmc1-E157D*, green – *dmc1-E157D rad51*, yellow – *dmc1-E157D mei5 rad51*. Strains used in experiments in the order in which they appear in figure, right to left and top to bottom: DKB3698, DKB6320, DKB6342, DKB6300, DKB3710, DKB6393, DKB6412.

### *dmc1-E157D rad51* exhibits a profound IH bias defect and a reduction in JM formation

We next examined Dmc1-E157D-mediated recombination in the absence of Rad51 using 2D gel electrophoresis. In a *rad51* mutant, Dmc1 carries out recombination, but there is a profound IH bias defect, and most recombination occurs between sisters [69]. The IH-dHJ/IS-dHJ ratio in *rad51* is 0.4 and the same ratio is displayed by *dmc1-E157D rad51* (Figures 4a,b). The defect in the IH/IS ratio is the result of increased IS-dHJs and decreased in IH-dHJs. The profound defect in IH bias in *dmc1-E157D rad51* contrasts with the *dmc1-E157D* single mutant, in which the IH-dHJ/IS-dHJ ratio is ∼1.6. We conclude that *rad51* is epistatic to *dmc1-E157D* with respect to its impact on partner choice. The impact of a *rad51* mutation on the IH/IS ratio in *dmc1-E157D* cells further supports the view that the mechanism of homolog bias is intact in *dmc1-E157D mei5* cells and therefore the conclusion that Mei5-Sae3 is not required for homolog bias. The levels of IS-dHJs, IH-dHJs, SEIs, and mcJMs are all reduced roughly 2-fold in *dmc1-E157D rad51* relative to *rad51* (Figure 4b); thus, the hyper-recombinant phenotype of *dmc1-E157D* cells is Rad51-dependent. These findings are also consistent with the observation that CO levels in *dmc1-E157D* are reduced about 2-fold by the *rad51* mutation (see Figure 2b).

**Figure 4.**
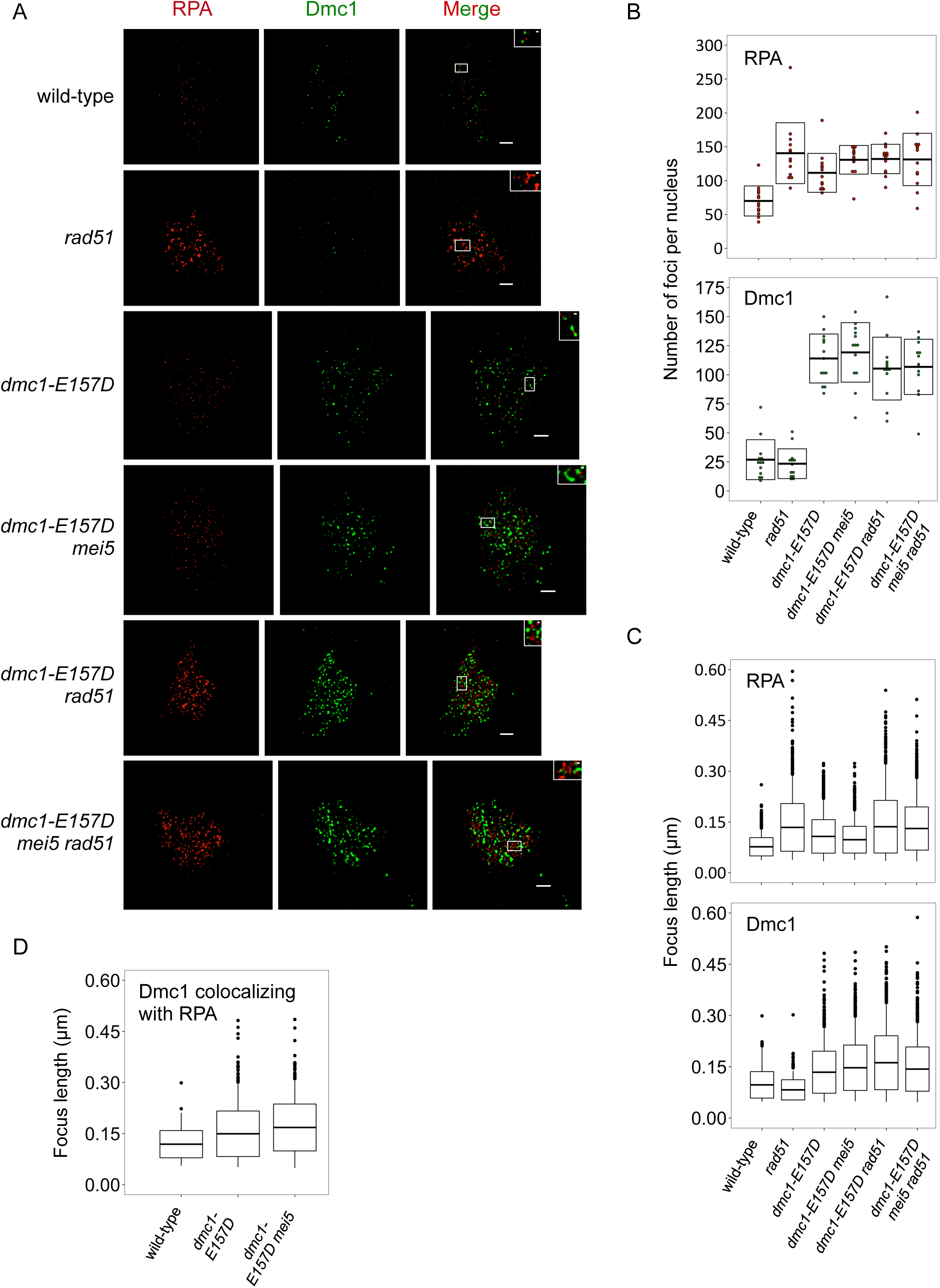
Super-resolution imaging shows abnormalities in RPA, Dmc1 foci in mutants. (a) Representative STED microscopy imaging of spread meiotic nuclei are shown for each strain. Scale bars represent 1 µm; scale bars in inset represent 0.1 µm. For *dmc1-E157D mei5*, time point was taken at 5 hours in sporulation media; for all other strains, time point was taken at 4.5 hours. (b) Quantitation of foci counts for Dmc1, RPA, is shown for each strain. For each strain, 13 randomly selected nuclei were quantitated. (c) Quantitation of RPA and Dmc1 foci lengths is shown for each strain. (d) Quantitation of Dmc1 foci lengths colocalizing with RPA is shown for the strains indicated. Strains used in this experiment in the order in which they appear in figure, top to bottom: DKB3698, DKB3710, DKB6342, DKB6300, DKB6393, DKB6412.

### JM formation is absent in triple mutant *dmc1-E157D mei5 rad51*

We also analyzed the *dmc1-E157D mei5 rad51* triple mutant by 2D gel electrophoresis. Surprisingly, while both *dmc1-E157D mei5* and *dmc1-E157D rad51* formed readily detectably levels of JMs (Figures 3a,c), and Dmc1 foci were detected in *dmc1-E157D mei5 rad51* spread meiotic nuclei (Figure 1c,d), no JMs were detected in the triple mutant (Figure 3c,d). Because *rad51* strains are genetically unstable, we constructed an independent *dmc1-E157D mei5 rad51* diploid and repeated this experiment to ensure that our original strain had not picked up an additional mutation that suppressed the formation of JMs. Meiotic JMs were also undetectable in the duplicate *dmc1-E157D mei5 rad51* strain (Supplemental Figures 3a,b). These 2D gel analyses are consistent with our findings that only ∼10% of cells progress through a meiotic division in *dmc1-E157D mei5 rad51*, and that there is hyper-resection and limited CO formation in this strain (Figures 2a,b). We conclude that recombination is further compromised in the triple mutant *dmc1-E157D mei5 rad51* than in either of the double mutants. These results provide additional evidence that although Dmc1-E157D’s activity is essentially Mei5-Sae3 independent in *RAD51^+^* cells, Mei5-Sae3 can promote limited Dmc1-E157D activity when Rad51 is absent.

### The defects associated with *dmc1-E157D* and *dmc1-E157D mei5* are independent of Rad51’s catalytic activity

One possible explanation for the results we obtained from our JM analysis is that the *dmc1-E157D* mutation changes the behavior of Dmc1 in a manner that activates the strand exchange activity of Rad51. This possibility is emphasized by previous results suggesting that Dmc1 itself inhibits Rad51’s strand exchange activity [79,80]. Normally, Rad51’s strand exchange activity is repressed by Dmc1 and by the meiosis-specific Rad51 inhibitor Hed1 [3,66]. However, it was important to determine if Rad51’s strand exchange activity plays a greater role in promoting recombination in *dmc1-E157D* cells than in wild-type [80]. To test this, we crossed the *rad51-II3A* mutation into our *dmc1-E157D* strains. The three alanine substitutions coded by *rad51-II3A* eliminate DNA binding site II, the secondary, low affinity DNA binding site required for homology searching. Rad51-II3A forms filaments, but lacks the ability to catalyze D-loop formation [3]. The results indicate that the *rad51-II3A* mutation does not alter the efficiency of JM formation in the *dmc1-E157D* mutant (Supplemental Figures 4a,b). This observation indicates that Dmc1, not Rad51, promotes the majority of homology search and strand exchange in *dmc1-E157D* cells, as is the case in wild-type cells. Thus, the hyper-recombinant phenotype observed in *dmc1-E157D* results from increased Dmc1 activity rather than activation of Rad51’s activity. On the other hand, *rad51-II3A* causes a greater reduction in spore viability in *a dmc1-E157D* background than in a wild-type background (Table 1, 17.0% and 82.9%, respectively, p < 0.01). The modest reduction in viability seen in *rad51-II3A* single mutants was previously interpreted to suggest that Rad51’s strand exchange is only required at a small subset of the roughly 200 DSB sites where Dmc1-dependent DSB repair fails [3]. In the context of this interpretation, the data presented here can be explained if the fraction of attempted recombination events that require Rad51’s strand exchange activity, although still small, is higher in *dmc1-E157D* than that in wild-type.

### Meiotic two-hybrid analysis indicates that direct Rad51-Dmc1 interaction is independent of Mei5

The results presented in Figure 3 show that Rad51 can impact Dmc1’s activity in the absence of Mei5-Sae3. To determine if Rad51’s influence on Dmc1 can be explained by direct interaction of the two proteins, we carried out meiotic two-hybrid analysis. A previous two-hybrid study in budding yeast using the conventional mitotic method detected a low level of direct interaction between Rad51 and Dmc1, although the authors of that study did not ascribe significance to the interaction because it was much weaker than that observed for homotypic Rad51-Rad51 and Dmc1-Dmc1 interactions [81]. We wished to determine if Mei5-Sae3 enhanced the interaction between the two proteins and therefore used the meiotic two-hybrid method to test the interaction in a cell type that expresses the accessory protein. As in the previous study, the level of interaction observed for Rad51-Dmc1 was much lower than that in the Rad51-Rad51 and Dmc1-Dmc1 homotypic controls, but nonetheless reproducibly higher than the background level observed in empty vector controls (Supplemental Figure 5). Importantly an equivalent two-hybrid signal was detected in a *mei5* null background as in a wild-type background indicating that, in this system, Rad51-Dmc1 interaction is independent of Mei5-Sae3.

### Super-resolution imaging of *dmc1-E157D* mutants reveals abnormalities in Dmc1 and RPA foci

Because Dmc1-E157D forms foci at high density, we expected that the wide-field microscopy method was not resolving closely spaced foci. Therefore, in order to obtain more accurate focus measurements, we re-examined chromosome spreads using STED microscopy, which improves the resolution limit from around 200 nanometers (nm) to under 50 nm (see Methods Section, Supplemental Figure 6a). For each strain, we imaged at least 13 randomly selected RPA-positive nuclei. The average number of RPA foci detected was lowest in wild-type (70.0 ± 22.2 foci). All other strains displayed higher average focus counts including *rad51* (140.5 ± 44.9 foci), *dmc1-E157D* (111.5 ± 28.8 foci), *dmc1-E157D mei5* (130.8 ± 21.2 foci), *dmc1-E157D rad51* (132.0 ± 21.7 foci), and *dmc1-E157D mei5 rad51* (131.3 ± 38.6 foci) (Figure 4b). We also measured focus lengths (Figure 4c), and found that wild-type RPA foci are the shortest (76.8 ± 27.0 nm), while *rad51*, *dmc1-E157D rad51*, and *dmc1-E157D mei5 rad51* foci are all significantly longer (134.0 ± 70.4 nm, 136.0 ± 77.8 nm, 130.8 ± 63.8 nm respectively; p < 0.01, Wilcoxon test), but not significantly different from one another (pairwise p = 0.53, 0.60, and 0.94, respectively). The fact that RPA foci are longer in these strains is unsurprising given that we observed hyper-resection in all of these strains by one-dimensional gel electrophoresis (Figure 2a). *dmc1-E157D* and *dmc1-E157D mei5* mutant RPA foci are significantly different from both wild-type and *rad51* mutants (107.4 ± 49.5 nm, 97.7 ± 39.6 nm respectively; p < 0.01), being an intermediate average length between the two.

The average number of Dmc1 foci per nucleus was similar in wild-type and *rad51* single mutants (26.9 ± 17.7 foci and 23.3 ± 12.8 foci, respectively, Figure 4b). All *dmc1-E157D* strains displayed higher than normal focus counts including *dmc1-E157D* (114.0 ± 21.0 foci), *dmc1-E157D mei5* (119.2 ± 25.6 foci), *dmc1-E157D rad51* (105.3 ± 27.0 foci), and *dmc1-E157D mei5 rad51* (106.8 ± 23.8 foci) (Figure 4b). This result is expected given that Dmc1-E157D forms numerous brightly staining foci in the absence of DSBs, whereas wild-type Dmc1 does not (Figure 1e). We also measured the lengths of these Dmc1 foci, and found that Dmc1 foci are significantly shorter in *rad51* (82.5 ± 30.0 nm, p < 0.01, Wilcoxon test) than wild-type (97.1 ± 38.8 nm) (Figure 4c), consistent with previous wide-field microscopy analyses [68]. Dmc1 foci are longer in all *dmc1-E157D* strains, including *dmc1-E157D* (134.1 ± 61.5 nm, p < 0.01), *dmc1-E157D mei5* (147.1 ± 66.4 nm, p < 0.01), *dmc1-E157D rad51* (161.9 ± 78.9, p < 0.01), and *dmc1-E157D mei5 rad51* (143.3 ± 64.7 nm, p < 0.01) relative to wild-type (Figure 4c).

Although measurements of Dmc1 focus lengths shows that Dmc1-E157D makes longer than normal filaments overall, the fact that the protein likely forms high levels of off-pathway foci in addition to forming foci at sites of recombination raises the possibility that the long filaments observed might only be off-pathway forms, with no appreciable change in the average length of recombinogenic filaments. Furthermore, the fraction of recombinogenic foci could differ in different strains. For example, off-pathway Dmc1 foci a larger fraction of the total in *dmc1-E157D* strains than in wild-type and *rad51*. To provide evidence that recombinogenic foci are longer on average, we examined the lengths of Dmc1 foci that colocalized with RPA. Given that all of the mutants have more RPA foci and some have more Dmc1 foci (Figure 4b), the level of fortuitous colocalization is expected to be higher in the mutants than in wild-type. We therefore estimated the frequency of fortuitous colocalization in all strains by a previously described method [74]. This method may yield an overestimate because the most focus dense region of each nucleus was used in the analysis. We eliminated any nuclei from our analysis if the level colocalization observed in the experimental image did not exceed the estimated frequency of fortuitous colocalization by more than 5%. Because both RPA and Dmc1 foci are more numerous in *dmc1-E157D rad51* and *dmc1-E157D mei5 rad51* (Figure 4b), and because both RPA and Dmc1 foci are on average larger in these strains (Figure 4c), the density of foci is much higher, and we were unable to identify a subset of Dmc1 foci in these strains that unambiguously colocalize with RPA (90.2% experimental and 91.4% fortuitous colocalization in *dmc1-E157D rad51*; 81.1% true and 80.0% fortuitous colocalization in *dmc1-E157D mei5 rad51*). In contrast, 10/13 nuclei wild-type nuclei (35.5% experimental and 18.8% fortuitous), 10/13 *dmc1-E157D* nuclei (70.1% experimental and 58.6% fortuitous colocalization), and 6/13 *dmc1-E157D mei5* nuclei (69.1% experimental and 57.1% fortuitous colocalization) met our criteria for analysis, indicating that the RPA-colocalization provides a meaningful criterion to identify a subset of Dmc1 foci enriched for recombinogenic as opposed to off-pathway structures. The average contour length of Dmc1 filaments that colocalized with RPA was 118.9 ± 40.0 nm in wild-type, or ∼100 nucleotides, similar to the corresponding value obtained using dSTORM, a different super-resolution light microscopy method [26]. The average focus length for RPA colocalizing Dmc1 foci in *dmc1-E157D* was significantly longer than in wild-type (149.5 ± 66.8 nm, or ∼ 160 nucleotides, Figure 4d, p<0.01, Wilcoxon test), and different from the total Dmc1 foci lengths in *dmc1-E157D* cells (134.1 ± 61.5 nm, Figure 4c). The average focus length for RPA colocalizing Dmc1 foci in *dmc1-E157D mei5* was also significantly longer than in wild-type (168.0 ± 66.4 nm, or ∼190 nucleotides, p <0.01), and different from the total Dmc1 foci lengths in that background (147.1 ± 66.4 nm, Figure 4c). This finding indicates that not only does Dmc1-E157D make longer foci overall, in *dmc1-E157D* and *dmc1-E157D mei5*, where we observe the hyper-recombinant phenotype, Dmc1 filaments associated with RPA are significantly longer than wild-type.

### Rhd54 promotes meiotic progression in *dmc1-E157D* cells

The cytological results presented above suggest that Dmc1-E157D is more likely than Dmc1-WT to form off-pathway filaments on dsDNA. DSB-independent foci are only easily detected for Dmc1-WT when Rdh54, the key translocase involved in disassembling them, is absent [41]. This observation suggested that Dmc1-E157D might be more resistant to dsDNA dissociation by Rdh54. To determine whether Rdh54 was active in *dmc1-E157D* mutants, we constructed the *dmc1-E157D rdh54* double mutant. If Rdh54 is inefficient at promoting Dmc1-E157D dissociation from dsDNA, loss of Rdh54 in the *dmc1-E157D* background should be inconsequential. Instead, we find that although both *dmc1-E157D* and *rdh54* single mutants progress through meiosis to form tetrads in which roughly 50% of spores are viable, the *dmc1-E157D rdh54* double mutant arrested in prophase and failed to form spores (Table 1; Supplemental Figure 7). Thus, Rdh54 is active in *dmc1-E157D* cells.

### Mei5-Sae3 is not required for the DSB-independent foci formed by Dmc1-WT protein in the absence of Rdh54

Dmc1-E157D differs from Dmc1-WT in that it forms high levels of off-pathway foci and does so independently of Mei5-Sae3. This suggests that although the mutant bypasses the requirement for Mei5-Sae3 with respect for forming recombinogenic foci, it might not fully recapitulate Mei5-Sae3 function because Mei5-Sae3’s has only been shown to display DSB-dependent foci; it was not known if Mei5-Sae3 is also required for the off-pathway Dmc1 complexes that accumulate when disassembly of dsDNA bound structures is blocked by an *rdh54* mutation. Therefore, to determine if Mei5-Sae3 is normally required for Dmc1 to form nascent complexes on dsDNA *in vivo*, we compared Dmc1 focus formation in *spo11 rdh54 mei5* to that in the *spo11 rdh54* double mutant; a *spo11* single mutant served as negative control. The controls generated the expected results with *spo11 rdh54* nuclei displaying an average of 37±14 Dmc1 foci/nucleus and *spo11* nuclei an average of only 3±4 foci/nucleus (Supplemental Figure 8). The *spo11 rdh54 mei5* triple mutant displayed an average of 37±13 foci, like the positive control, indicating that focus formation in *spo11 rdh54* is Mei5 independent. Thus, a key component of Mei5-Sae3 function appears to be specific to promoting filaments on ssDNA. Dmc1-E157D appears to bypass the requirement for Mei5-Sae3 for filament formation on ssDNA, but does so without displaying the ssDNA-specific function normally provided by Mei5-Sae3.

## Discussion

### The mechanism of Mei5-Sae3-mediated Dmc1 filament formation

Dmc1-E157D was designed to mimic RecA-E96D. The RecA-E96D mutation shortens the length of a critical amino acid side chain in the ATPase active site, increasing the distance between the water molecule that acts as the nucleophile for hydrolysis, and the activating carboxylate [71]. The mutation dramatically reduces that rate of ATP hydrolysis thereby maintaining RecA in the ATP-bound form, which is active for DNA binding, homology search, and strand exchange. Due to the high sequence conservation of this site, Dmc1-E157D is very likely to be defective in ATPase activity, like RecA-E96D.

Assuming this prediction is correct, our results provide *in vivo* support for the conclusion of Chi and colleagues that Swi5-Sfr1 acts to stabilize Rad51 filaments by promoting ADP release, thereby maintaining the filament in the active, ATP-bound form [59]; a mutation designed to favor the ATP bound form of Dmc1 bypasses the normal requirement for Mei5-Sae3. On the other hand, the regulatory defects observed in Dmc1-E157D suggest that the function of Mei5-Sae3-mediated regulation involves more than overall enhancement of Dmc1 filament stability, because the Dmc1-E157D mutant displays abnormally high levels of *spo11*-independent Dmc1-E157D binding to chromosomes (Figure 1e). This finding suggests that stabilizing the ATP-bound form of Dmc1 alone is insufficient to account for the mechanism of Mei5-Sae3 function. Supporting this view, we find that although Mei5-Sae3 is required for cytologically detectable Dmc1 focus formation at sites of DSBs in wild-type cells, it is not required to observe the off-pathway dsDNA-bound foci formed on dsDNA by Dmc1-WT in Rdh54 deficient cells (Supplemental Figure 8). This interpretation is consistent with prior observation of direct binding of Mei5-Sae3 to the ssDNA-specific binding protein RPA as well as the ability of Mei5-Sae3 to enhance Dmc1 activity in the presence of RPA [50,57,60]. Thus, Mei5-Sae3 appears to combine the ability to enhance Dmc1 filament stability with the ability to specifically promote filament formation on ssDNA rather than dsDNA.

The ability of Dmc1-E157D to form functional filaments on ssDNA *in vivo* in the absence of Mei5-Sae3, and to do so by a mechanism involving filament stabilization, opens the possibility that recruitment/nucleation of Dmc1 filaments on RPA coated ssDNA in normal cells is independent of Mei5-Sae3. Given that Mei5-Sae3 binds directly to both Dmc1 and RPA [52,53,60], we continue to favor models in which Mei5-Sae3 plays a role in recruitment/nucleation of Dmc1 filaments. We note, however, that Dmc1 could be recruited to sites of DSBs through its interactions with RPA [50], and that nucleation, but not filament elongation, could be Mei5-Sae3 independent. Dmc1 nucleation events might be undetected in the absence of Mei5-Sae3 because the resulting filaments never elongate to lengths sufficient to reach the threshold of cytological detection. It is also possible that Rad51 is normally partially responsible for Dmc1 recruitment/nucleation, in addition to its roles in filament stabilization and homolog bias. These considerations highlight the need for further studies on the mechanism of Dmc1 recruitment/nucleation on RPA coated ssDNA tracts *in vivo*.

### The role of Rad51 in Dmc1 filament dynamics

The absence of foci observed in *mei5*, *sae3*, and *mei5 sae3* mutants, and the dimmer foci observed in *rad51* mutants, indicates that normal Dmc1 nucleoprotein filament formation involves both proteins. The fact that recombination and DSB-dependent focus formation in *rad51* yeast depends on Mei5-Sae3 suggests that Mei5-Sae3 is epistatic to Rad51. Furthermore, formation of brightly staining Mei5-Sae3 foci depends on Rad51, as does formation of brightly staining Dmc1 foci [52,68]. These dependency relationships raised the possibility that Rad51’s ability to influence Dmc1 filaments might require a direct interaction between Rad51 and Mei5-Sae3 [82]. However, the data presented here indicate that Rad51 promotes formation of functional Dmc1 filaments on ssDNA independently of Mei5-Sae3, thus Rad51’s normal influence on Dmc1 filament dynamics does not require, and may not involve, Mei5-Sae3 binding to Rad51.

Our data clearly demonstrate that *dmc1-E157D* functions independent of Mei5-Sae3, yet the mutant is more dependent on Rad51 than the wild-type protein. Whereas *dmc1-E157D mei5* forms COs at a level nearly equivalent to wild-type, *dmc1-E157D rad51* suffers a dramatic reduction in CO formation, and experiences hyper-resection (Figures 2b,4c). In addition, 2D gel electrophoresis shows that JM formation in *dmc1-E157D mei5* is equivalent to *dmc1-E157D*, while the JMs formed in the *dmc1-E157D rad51* background are significantly reduced relative to *dmc1-E157D,* and show an IH bias defect like the *rad51* single mutant (Figures 3b, 3d). Thus, a mutation that alleviates the need for one accessory factor, Mei5-Sae3, makes Dmc1 more dependent on a second accessory factor, Rad51. This finding provides further evidence that Mei5-Sae3 and Rad51 functions are not interdependent with respect to enhancing the formation of functional Dmc1 filaments. If this were the case, a mutation that bypasses the requirement for one factor would also bypass the requirement for the second factor. This model accounts for the partial dependency of Mei5-Sae3 foci on Rad51; the reduction of Mei5-Sae3 focus intensity observed in *rad51* mutants is expected if Dmc1 filaments are bound along their lengths by Mei5-Sae3, and loss of Rad51 results in shorter Dmc1 filaments.

Rad51 is likely to impact Dmc1 filament dynamics by direct interaction. Although a previous study did not ascribe significance to the low level of interaction detected between budding yeast Rad51 and Dmc1 [81], two-hybrid studies in other organisms detected significant levels of Rad51-Dmc1 interaction, albeit at low levels compared to homotypic interactions [83–85]. Budding yeast Rad51 binds Dmc1 directly when pure proteins are mixed [50], consistent with similar observations in other organisms [83–85]. Using the meiotic two-hybrid method, we were able to detect Rad51-Dmc1 interaction during meiotic prophase of budding yeast, and to show that this interaction does not depend on Mei5-Sae3. These findings provide additional evidence that Rad51 and Mei5-Sae3 influence Dmc1 DNA binding dynamics independently. The finding that Rad51-Dmc1 interaction occurs, but is weaker than homotypic interactions, is consistent with a single molecule study that showed mixtures of Rad51 and Dmc1 form predominantly homo-filaments on DNA [21], and with prior cytological studies that showed the foci formed by Rad51 and Dmc1 lie adjacent to one another rather than being perfectly colocalized [51,81,86]. Finally, we note that direct interaction between the two proteins can account for the observation that Rad51 can stimulate Dmc1-mediated D-loop formation in the absence of other proteins [3].

How might Mei5-Sae3 and Rad51 promote filament stability by independent mechanisms? There are at least two basic mechanisms that could contribute to filament stability. First, an accessory protein could promote the high-affinity ssDNA binding form. Second, if a strand exchange protein is normally subject to enzymatically-driven disassembly, an accessory protein might act by specifically blocking the activity of that enzyme. Mei5-Sae3’s role in filament stabilization *in vivo* almost certainly involves direct enhancement of DNA binding activity during nucleation and/or elongation, as is the case for Mei5-Sae3 homolog Sfr1-Swi5 [62]. Rad51 might also enhance binding directly, by reducing the off-rate of protomers from filaments. For example, a Rad51 monomer bound to the end of a Dmc1 filament might drastically reduce the off-rate of the adjacent Dmc1 protomer with a strong overall effect on filament stability, given that disassembly of filaments is expected to occur from filament ends [87].

Alternatively, Rad51 may block a mechanism that actively dissociates Dmc1 filaments. Although no active assembly mechanism has been identified for Dmc1 filaments, active disassembly could involve a helicase mechanism, similar to that mediated by UvrD and Srs2 [16–19]. One observation that appears to be at odds with the idea that Rad51 functions by blocking an Srs2-like mechanism is that Rad51 can stimulate Dmc1’s D-loop activity in a purified system that does not include an ssDNA-specific helicase. However, it is possible that the *in vitro* activity of Rad51 in stimulating Dmc1 does not fully recapitulate the *in vivo* function of the protein. This possibility is emphasized by previous work on the Rad51 accessory protein Rad55-Rad57. Both subunits of the Rad55-Rad57 heterodimer are structurally similar to Rad51. Rad55-Rad57 stimulates Rad51 activity *in vitro*, but *in vivo* it functions to limit the Rad51 strippase activity of Srs2 [88,89]. Thus, Rad51’s impact on Dmc1 activity *in vitro* might similarly not fully represent its *in vivo* role in promoting stable Dmc1 filaments.

A model invoking inhibition of Dmc1-ssDNA filament disassembly can account for the fact that *dmc1-E157D rad51* forms fewer JMs relative to *DMC1^+^ rad51* (Figure 3d). Like Dmc1-E157D, the Rad51 ATPase mutant Rad51-K191R is defective in recruitment to DSB-associated tracts of ssDNA *in vivo*. The DNA binding defect of Rad51-K191R is partially suppressed by deletion of *SRS2* or by overexpression of *RAD54* [45,46]. These findings suggest that the recruitment defect displayed by Rad51-K191R results from a combination of the protein’s DNA binding defect, increased off-pathway dsDNA binding, and active disassembly of the Rad51-K191R filaments that do form at DSB-associated tracts of ssDNA [47].

If Dmc1-E157D filaments form more slowly than wild-type filaments as a result of increased off-pathway binding and thus a decreased pool of free Dmc1 protomers, Dmc1-E157D filaments may be acutely sensitive to disassembly and/or end dissociation, thus both models can explain Dmc1-E157D’s increased dependency on Rad51. In addition, these models can account for the more severe phenotype of the *dmc1-E157D mei5 rad51* triple mutant compared to the *dmc1-E157D rad51* double mutant as a consequence of Mei5-Sae3 having a limited ability to block dissociation, or being able to promote fast reassembly. Such an activity of Mei5-Sae3 might be inconsequential for Dmc1-E157D-DNA binding dynamics *in vivo* when Rad51 is present, explaining why the phenotypes of *dmc1-E157D* and *dmc1-E157D mei5* are nearly identical.

### Mei5-Sae3 is not required for IH bias

The results presented here also reveal for the first time that although both Rad51 and Mei5-Sae3 promote the formation of stable Dmc1 filaments, Mei5-Sae3 differs from Rad51 in that Mei5-Sae3 is <u>not</u> required for homolog bias while Rad51’s function is. This conclusion could not have been arrived at based on earlier observations because recombination is blocked prior to formation of joint molecules in *mei5 DMC1^+^* and *sae3 DMC1^+^* cells; bypass of the requirement for Mei5-Sae3 for formation of functional filaments allowed us to assess the role of Mei5-Sae3 during choice of recombination partner at the D-loop formation stage. Previous work showed that Rad51 and Dmc1 are both required for homolog bias [69,80]. The results here show that the cooperation between Rad51 and Dmc1 required for IH bias involves a Rad51-dependent mechanism that is independent of Mei5-Sae3. This interpretation is consistent with the fact that, in other species, homologs of Mei5-Sae3 regulate Rad51 activity, suggesting that the Mei5-Sae3 family of accessory proteins solves a problem common to both Rad51 and Dmc1, and not unique to meiotic recombination.

Chromatin immunoprecipitation experiments have shown that cells lacking both Rdh54 and Rad54 fail to recruit Dmc1 to DSB hotspots as a consequence of sequestration caused by off pathway DNA binding. The failure to recruit Dmc1 to tracts of ssDNA accounts for the hyper-resection seen in *rad54 rdh54* double mutants [41,90]. Given that Dmc1-E157D forms foci in the absence of DSBs, and that it is modeled on RecA-E96D, which displays a lower than normal off-rate for dsDNA binding, one might expect that Dmc1-E157D is less efficiently removed from dsDNA by Rdh54 (and Rad54). Surprisingly, we find no evidence for a decrease in CO formation or for hyper-resection in *dmc1-E157D* (Figures 2a,b). Moreover, there is no accumulation of SEIs, which might be expected if Rdh54 were unable to remove Dmc1 from the 3’ end of the heteroduplex DNA to allow for recombination-associated DNA synthesis (Figures 3a,b). We also find that the high spore viability and meiotic progression observed in *dmc1-E157D* mutants is strongly dependent of Rdh54, indicating that Rdh54 is active in *dmc1-E157D* mutants (Table 1, Supplemental Figure 4). Thus, although Dmc1-E157D forms more off-pathway filaments than Dmc1-WT, Rdh54 appears to be capable of dissociating them.

### Dmc1-E157D forms abnormally long filaments and is hyper-recombinant for certain recombination events

Although levels of IH CO intermediates and products are similar to those in wild-type, *dmc1-E157D* and *dmc1-E157D mei5* display higher than normal levels of certain types of recombination intermediates and products including IS-dHJs, mcJMs, and ectopic COs. For simplicity, we will refer to these unusual types of recombination events collectively as “aberrant,” but we emphasize that all three types are observed at low levels in wild-type. IS-dHJs, mcJMs, and ectopic COs are all elevated about 3-fold in *dmc1-E157D* and *dmc1-E157D mei5* cells (Figures 2b,3b,3d). The combination of aberrant recombination phenotypes observed in *dmc1-E157D* cells is reminiscent of that reported for *sgs1*, *top3*, and *rmi1* mutants during meiosis [91–93]. Sgs1, Top3, and Rmi1 have been shown to form a complex, STR, that disassembles D-loops [94–96]. In addition, during mitotic recombination, STR was shown to have a role in disassembling aberrant invasion events in which a single Rad51 filament invades two or more donor molecules (“multi-invasions”, or MIs) [97]. This role of STR in MI disassembly was proposed to account for at least some of the phenotypes observed in the absence of Sgs1, Top3, or Rmi1 during meiosis [93]. In this context, maturation of a MI into a mcJM, followed by resolution of the MI, can account for the increase in mcJMs, IS-dHJs, and ectopic recombination observed in these mutants [98]. Further evidence that MIs account for the meiotic STR mutant phenotypes is the fact that both MIs and JMs in the *sgs1*, *top3*, or *rmi1* mutant backgrounds are highly dependent on structure-selective nucleases Mus81-Mms4, Slx1-Slx4, and Yen1 [92,93,97,99–101].

Two possibilities account for why *dmc1-E157D* and *dmc1-E157D mei5* are phenotypically similar to STR mutants. Dmc1-E157D may form the same number of aberrant intermediates as wild-type, but STR-mediated disassembly could be rendered less efficient as a consequence of enhanced binding activity of Dmc1-E157D compared to Dmc1-WT. Arguing against this possibility is the fact that there is no increase in SEIs in *dmc1-E157D* and *dmc1-E157D mei5* cells compared to wild-type (Figures 3b, 3d), which is expected if the mutant protein prevents nascent D-loop disruption. Moreover, we showed that Rdh54 promotes meiotic progression in *dmc1-E157D* (Table 1, Supplemental Figure 7), implying that Rdh54 is competent to remove Dmc1-E157D from dsDNA.

An alternative model to account for the defects associated with *dmc1-E157D* and *dmc1-E157D mei5* is that Dmc1-E157D makes more aberrant D-loops than Dmc1-WT. In this model, STR, and possibly other helicases, disassemble aberrant D-loops as normal, but the mutant protein generates more MIs than Dmc1-WT. The two regions of homology engaged in such MI events could be on one sister and one homolog, or on both of the homologs, likely engaging one template at the allelic site, and one at the ectopic site. The formation of the MIs can account for the increased mcJMs, while processing of MIs to yield fully repaired chromatids can explain the increases in IS-dHJs and ectopic COs [98]. Drawing on the “intersegmental contact sampling” model of homology search [102], we propose Dmc1-E157D makes more MIs as a consequence of making longer filaments (Figure 5). The intersegmental contact sampling model maintains that a filament has a polyvalent interaction surface capable of simultaneously searching multiple, non-contiguous DNA regions for homology [102]. Longer filaments are able to search duplex DNA more efficiently, as a consequence of being able to engage in a greater number of simultaneous interactions. We have demonstrated that Dmc1-E157D forms longer filaments *in vivo* (Figure 4c). We posit that because filaments are longer, Dmc1-E157D engages in a higher number of simultaneous searching interactions that results in more frequent homology-dependent engagement of two different regions of homology by a single filament. In addition, though these aberrant recombination events are increased in *dmc1-E157D*, they also make up a substantial fraction of the recombination events observed in wild-type [91,97]. Consistent with this finding, 14% of wild-type Dmc1 foci that colocalized with RPA or ∼1.2 foci/nucleus were longer than 149 nanometers in length, the average length of Dmc1-E157D foci that colocalize with RPA in *dmc1-E157D* (Figure 4d). This finding suggests that although most foci are much shorter than 149 nanometers in wild-type, long filaments do occasionally form. Supporting the proposal that longer than normal filaments are responsible for higher than normal levels of MIs, previous work showed that (1) if longer ssDNA substrates are used, there is a higher incidence of MI formation [103]; and (2) Rad55-Rad57 promotes both longer Rad51 filaments and the formation of MIs [89,98].

**Figure 5.**
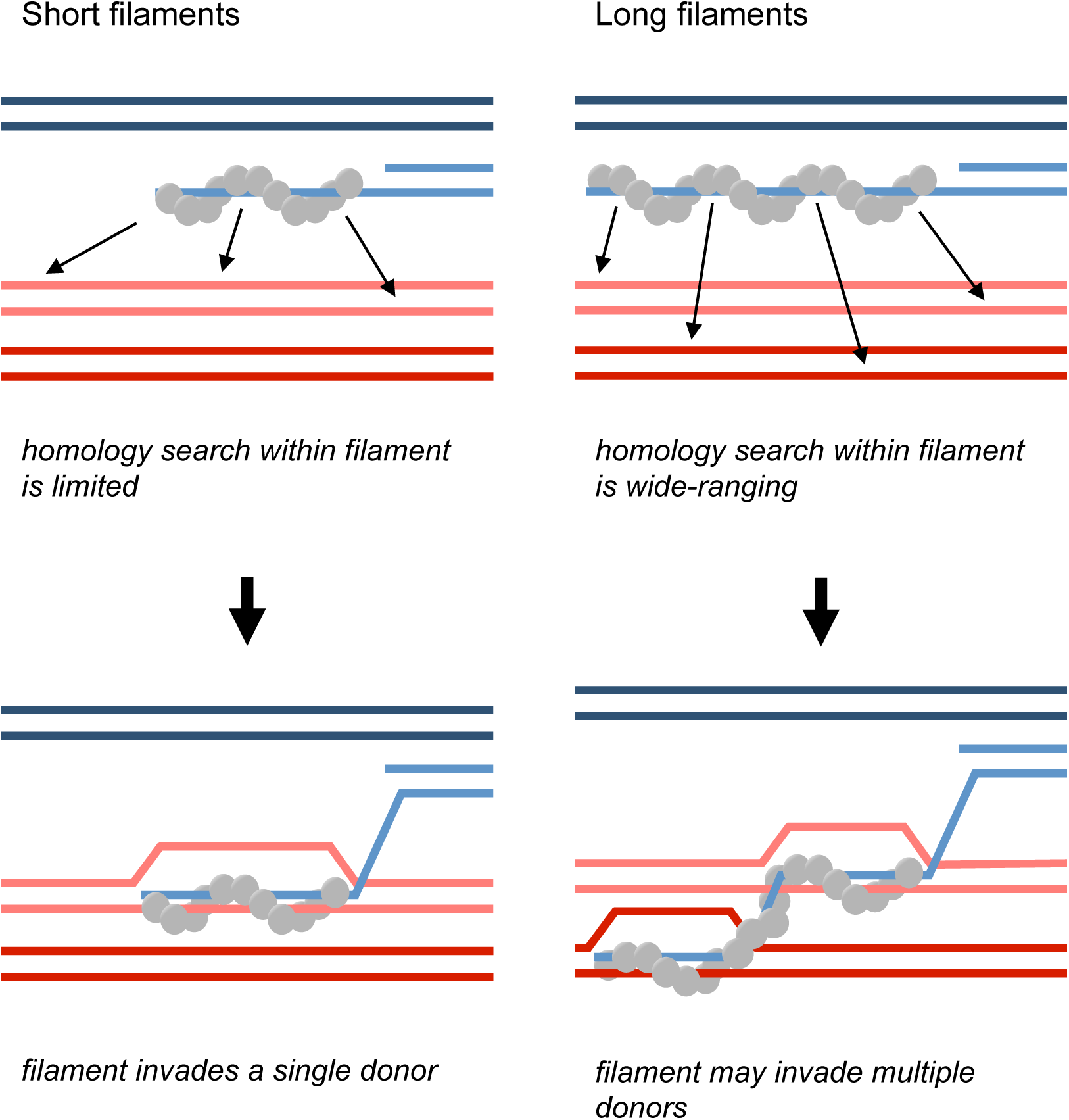
Model for regulation of filament length *in vivo*.

The aberrant event hyper-recombinant phenotype displayed by Dmc1-E157D is Rad51-dependent. The mechanism responsible for Rad51’s role in promoting the aberrant hyper-recombinant activity of Dmc1-E157D remains to be determined. Analysis of RPA co-localized foci provided evidence that Dmc1-E157D forms longer filaments on ssDNA in otherwise wild-type cells and in *mei5* single mutants. The mutant protein also forms long filaments on dsDNA, given that long filaments are observed in *spo11* mutants. Because both RPA and Dmc1 foci counts are increased in *dmc1-E157D rad51* and *dmc1-E157D mei5 rad51* mutants (Figure 4b), and because both RPA and Dmc1 foci are also larger in these mutants (Figure 4c), it was not possible to identify a sub-population that we could be confident was enriched for ssDNA bound structures in these mutants. As a result, it is unclear if the dependency of Dmc1-E157D’s hyper-recombinant phenotype on Rad51 reflects a requirement for Rad51 in forming long Dmc1 filaments on ssDNA, or if Rad51 plays some other role in promoting the high level of aberrant recombination events observed in Dmc1-E157D. It is clear, however, that Rad51’s homology search and strand exchange activities are not required for the aberrant hyper-recombinant phenotype observed in *dmc1-E157D* cells because the *rad51-II3A* mutation had no impact on the phenotype.

We speculate that the lengths of RecA-family strand exchange filaments are limited by regulatory mechanisms that evolved to prevent homology-dependent translocations and other genome rearrangements. Limiting filament lengths may limit the ability of filaments to simultaneously engage more than one homologous target sequence. In this regard, it is relevant that the single molecule study that provided evidence for intersegmental transfer did not detect any homology-dependent target engagement with the shortest ssDNA substrate examined, which was 162 nucleotides in length [102]. However, *in vivo*, Dmc1 filaments are typically ∼100 nucleotides in wild-type cells (Figure 4c) [26]. Thus, it is possible that the cost of MIs to genome stability has limited the length of strand exchange filaments such that intersegmental searching is limited or prevented *in vivo.* Alternatively, homology search may proceed by an intersegmental contact sampling mechanism, but filament lengths may nonetheless be limited to avoid genome-destabilizing MIs.

## Materials and Methods

### Yeast Strains

The yeast strains used in this study are listed in Supplemental Table 1. All yeast strains are isogenic derivatives of strain SK-1.

To construct the *dmc1* point mutants, DKB plasmid pNRB628 containing the *DMC1* open reading frame, a 701 base pair upstream homology arm, the *TEF1* promoter, the *natMX4* open reading frame, the *ADH1* terminator, and a 40 base pair downstream homology arm, was modified by Gibson assembly to include the desired point mutations. *dmc1::LEU2-URA3-KAN* haploid yeast (DKB129, DKB130) were transformed with a linear PCR fragment containing the homology arms, the mutated *dmc1* open reading frame, and the *natMX4* (for resistance to nourseothricin sulfate, or cloNAT) selectable marker. Yeast were outgrown in 5 milliliters liquid YPDA for 4.5 hours at 30°C in a culture rotator, then plated on selective media and allowed to grow at 30°C for 3 days. After 3 days, colonies were struck out on the selective media and on 5-fluoroorotic acid (5-FOA), which selects against *URA3^+^*yeast and therefore identifies clones that have lost the *dmc1::LEU2-URA3-KAN* allele. Those colonies that grew on the cloNAT media and did not grow on the 5-FOA plates were tested to confirm proper targeting by polymerase chain reaction, and then confirmed via sequencing.

### Meiotic Time Courses

Yeast cultures were induced to undergo synchronous meiosis as described previously [51]. Appropriate samples were collected at time points indicated in figures.

### Spore viability

Spore viability was determined by tetrad dissection as the percent of spores that germinate and form a colony on a YPDA plate relative to the number expected if all dissected spores had lived.

### Preparation and staining of spread yeast nuclei

Surface-spreading and immunostaining of meiotic yeast chromosomes on glass slides was performed as described previously [104]. Primary antibodies were used at the following dilutions: purified anti-goat Dmc1 bleed #4 DKB antibody #192 (1:800), anti-rabbit Rad51 bleed #2 DKB antibody #159 (1:1000), anti-rabbit RFA2 (1:1000), and anti-rabbit Hop2 bleed #3 DKB antibody #143 (1:1000). Secondary antibodies were used at a dilution of 1:1000 and included: Alexa Fluor 488 chicken anti-goat (Invitrogen by ThermoFisher Scientific), Alexa Fluor 594 donkey anti-rabbit (Invitrogen by ThermoFisher Scientific), Alexa Fluor 594 donkey anti-goat (Invitrogen by ThermoFisher Scientific) and Alexa Fluor 488 donkey anti-rabbit (Invitrogen by ThermoFisher Scientific). Images were collected on a Zeiss Axiovision 4.6 wide-field fluorescence microscope at 100X magnification. The same imaging parameters were used for all samples.

### Wide-field microscopy analysis

For each strain, 50 or more adjacent and randomly selected nuclei were imaged. A field of nuclei was chosen for analysis based on the DAPI staining pattern. Nuclei were scored as focus positive if there were 3 or more immunostaining foci in a given nucleus. Due to focus crowding in wide-field images, it was not possible to generate reliable focus counts using automated methods. Therefore, focus counts were determined by eye for the experiments reported in Supplemental Figure 8.

### One-dimensional gel electrophoresis

One-dimensional gel electrophoresis at the *HIS4:LEU2* meiotic hotspot was performed as follows. 15 milliliter sporulation media samples were collected at time points indicated from meiotic cultures. Sodium azide was added to a final concentration of 0.1%. Cells were spun down at 3000 rpm in tabletop clinical centrifuges for 5 minutes, then the supernatant was removed and the pellet was frozen at −20°C. DNA was then purified as described previously [105]. Approximately 2 micrograms DNA per sample was then digested with XhoI restriction enzyme (New England BioLabs) and processed as described previously [105]. Samples were then run on a 0.6% agarose gel at 2V/cm for 24 hours, followed by Southern blotting as described previously [78].

### Two-dimensional gel electrophoresis

Two-dimensional gel electrophoresis at the *HIS4:LEU2* meiotic hotspot was performed as previously described [3].

### Meiotic two-hybrid analysis

Analysis of Rad51-Dmc1 interaction in meiotic cells was performed using the meiotic two-hybrid method [106]. DNA binding domain constructs were transformed into *MATa* haploid strains DKB6431 (*MEI5^+^*) and DKB6429 (*mei5*) and activation domain constructs were transformed into *MATα* haploid strains DKB6430 (*MEI5^+^*) and DKB6428 (*mei5*). Independent transformants were mated to generate the diploid strains used for meiotic two hybrid experiments. 5 ml cultures were grown for 72 hours in synthetic tryptophan leucine dropout media to maintain 2µ plasmids and then transferred to YPD medium at OD600=0.2, and then grown for two generations before being transferred to SPS medium overnight, after which sporulation was induced on SPM-1/5COM medium. Recipes for media are as described previously [51]. Samples were prepared for β-galactosidase assays after 6 hours and 18 hours. The plasmids used for the two-hybrid studies were derived from pGAD-C1 [107] for activation domain fusions, and from pCA1 a gift from Scott Keeney [106] for DNA binding domain fusions. Note that this system uses *E. coli lexA* as DNA binding domain for hybrid constructs in combination with a *lex-op::lacZ* reporter construct [106]. Plasmid designations and the markers carried by the plasmids were as follows: Dmc1BD=pNRB729 *2µ, TRP1, P_DMC1_-DMC1-lexA, ampR, ori; Dmc1AD=pNRB271 2µ, LEU2, P_ADH_-GAL4-AD::DMC1, ampR, ori*; Rad51BD=pNRB727 *2µ, TRP1, P_DMC1_-lexA-Rad51, ampR, ori*; Rad51AD=pNRB688 *2µ, LEU2, P_ADH_-GAL4-AD::RAD51, ampR, ori*; ΔBD=pNRB728 *2µ, TRP1, P_DMC1_-lexA, ampR, ori*; and ΔAD=pNRB267 *2µ, LEU2, P_ADH_-GAL4, ampR, ori.* Plasmid sequences are available on request.

### Immunofluorescence imaging by stimulation depletion (STED) microscopy

Spreads were stained using a protocol described previously [104] with the following modifications. Spreads were dipped in 0.2% Photo-Flo (Kodak) for 30 seconds, the excess was tapped off, and then the slides were washed in 1X TBS for 5 minutes. Spreads were then blocked with 300µL 3% BSA in 1X TBS. Following blocking, spreads were incubated with anti-goat Dmc1 (1:800) and anti-Rabbit RPA (1:1000) for ≥16 hours at 4°C. Slides were then washed in 1X TBS + 0.05% Triton X-100 for 5 minutes with gentle rocking 7 times. Spreads were incubated with fluorochrome-conjugated secondary antibodies Alexa Fluor 594 donkey anti-goat and Alex Fluor 488 donkey anti-rabbit (1:1000) (ThermoFisher Scientific) for 2 hours at 4°C, followed by washes as described. Slides were allowed to dry completely in fume hood, then 35µL Vectashield (Vector Laboratories, Inc.) was added, a coverslip was placed atop the slide, and the coverslip was sealed with nail polish.

Imaging was conducted on a Leica SP8 3D, 3-color Stimulated Emission Depletion (STED) Laser Scanning Confocal Microscope at the University of Chicago Integrated Light Microscopy Core Facility. The same imaging parameters were used for all strains. Images were deconvolved using Huygens software and applying the same settings for each image. Resolution is reported based on measurements taken from deconvolved images.

### STED microscopy analysis

To quantitate the number of foci in each nucleus, the image channels were separated, and each channel image was converted to a binary image in ImageJ. The “Analyze Particles” function was used to obtain information regarding the number of foci in an image, the coordinates of the center of each focus, and the major length of each focus. The same settings were used to analyze all images. Colocalization between Dmc1 and RPA was scored in R using the coordinates given by ImageJ to calculate the distance between a given Dmc1 focus and all RPA foci in the nucleus. A Dmc1 focus was scored as colocalizing with a RPA focus if the nearest RPA focus was less than the length of that Dmc1 focus plus a preset RPA value that was calculated for each strain. The RPA value was calculated based on one half of the average length of all RPA foci in that sample plus one half of two standard deviations of that RPA length. This means that if a given Dmc1 focus is sitting side-by-side with an RPA focus, the distance between it and the center of the nearest RPA focus can be the length of that Dmc1 focus plus one half the average length of all RPA foci in that strain background, plus one half of two standard deviations of the RPA focus lengths. This calculation attempts to take into account the fact that both RPA foci and Dmc1 focus lengths vary from sample to sample. Plots and statistical tests were carried out in R using the ggplot and ggpubr packages.

### Meiotic whole cell lysate, SDS-PAGE, and Western blotting

4 milliliters of meiotic culture was collected at the appropriate time point. Tricholoroacetic acid was added to a final concentration of 10% weight/volume. Samples were placed in a 60°C water bath for 5 minutes, then placed on ice for 5 minutes. Next, samples were spun down at 3000 rotations per minute in a low-speed centrifuge, the supernatant removed by aspiration, and pellet then washed in ddH_2_O. The pellet was then re-suspended in 1X-SDS-PAGE (60 mM TrisHCl pH 6.8, 0.05% SDS, 100 mM DTT, 5% glycerol) buffer supplemented with 50 mM sodium PIPES pH 7.5 to the appropriate concentration according to the optical density of cells in the sample. The samples were then boiled for 10 minutes, spun down, and pellets stored at −20°C.

A 12% SDS-polyacrylamide gel was prepared, and 30 microliters of each sample was run at 120V for 1.5 hours alongside 20 nanograms purified Dmc1 protein. Samples were then transferred to Merck Millipore Limited Immobilon-P Transfer Membrane for 16 hours at 50V at 4°C. The membrane was then blotted using anti-goat Dmc1 (1:1000) primary antibody and an anti-goat HRP-conjugated secondary antibody (1:1000).

## Acknowledgments

We thank Wolf-Dietrich Heyer for helpful discussions and suggestions. Thanks to Akira Shinohara for the gift of the anti-rabbit RFA2 antibody, and Scott Keeney for the gift of the meiotic two-hybrid strains. We are grateful to Vytas Bindokas and Christine Labno for assistance with STED microscopy. Melissa Castiglione constructed the strains and one of the plasmids used in the two-hybrid analysis. This work was supported by NIGMS grant GM50936 and NCATS grant 1UL1TR002389-01 to DKB. DFR was partly supported by the NIH Genetics & Regulation Training Grant (T32 GM07197).

## Author Contributions

DFR and DKB jointly conceived the project and planned the experiments. DFR collected and analyzed the data shown in Figures 1-5, Supplemental Figures 1-4 and 6. JTG collected and analyzed the data for Supplemental Figure 5. DKB collected and analyzed the data for Supplemental Figures 7 and 8. DFR wrote the original manuscript. DFR and DKB jointly revised the manuscript. DKB was responsible for funding acquisition and project administration.

---

**Supplemental Figure 1.**
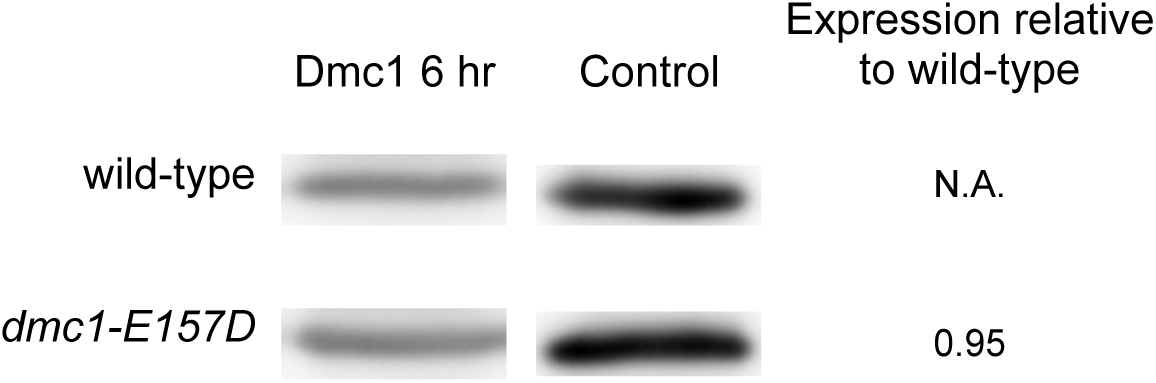
*DMC1* expression for wild-type, *dmc1-E157D*. Left column, W. blot against Dmc1 for 5µL sample prepared from meiotic yeast cultures at 6 hours as described in Methods Section for each strain. Control column is 20 ng purified Dmc1 protein that was run in parallel with sample and used to quantitate blots. Sample concentration is estimated concentration in comparison to 20 ng purified Dmc1 protein. Similar results were obtained from an independent meiotic time course. Strains used in this experiment in the order in which they appear in figure, top to bottom: DKB3698, DKB6342.

**Supplemental Figure 2.**
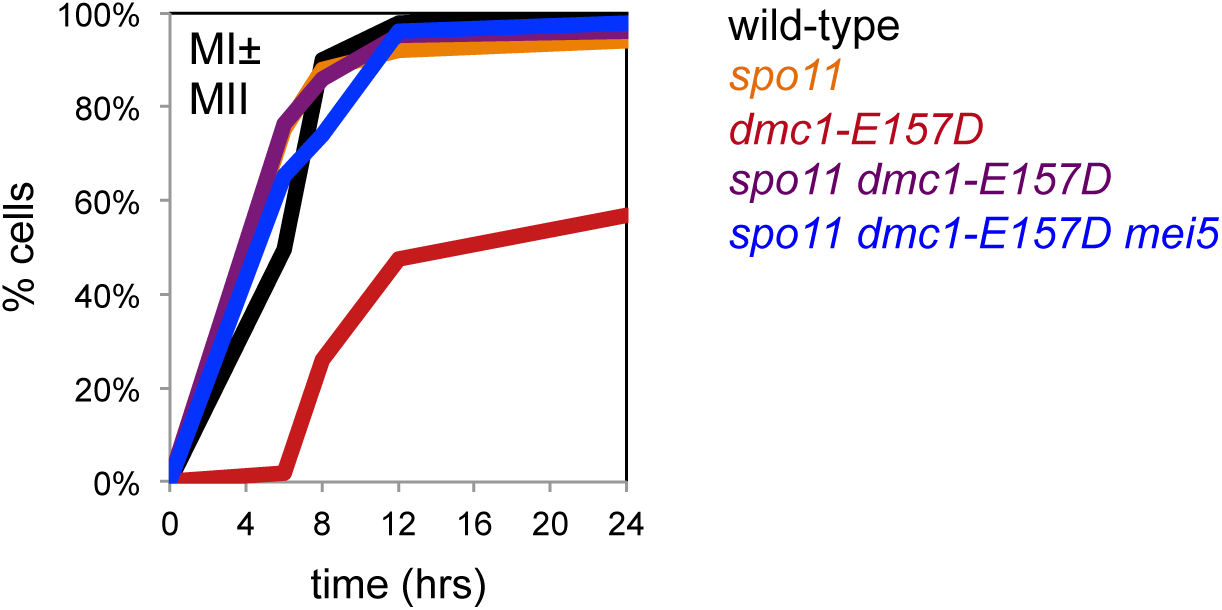
*spo11* suppresses the meiotic progression defect associated with *dmc1-E157D*. Meiotic progression data for strains indicated. For each time point, ≥50 cells were scored. Strains used in this experiment in the order in which they appear in figure, top to bottom: DKB3698, DKB2123, DKB6342, DKB6419, DKB6425.

**Supplemental Figure 3.**
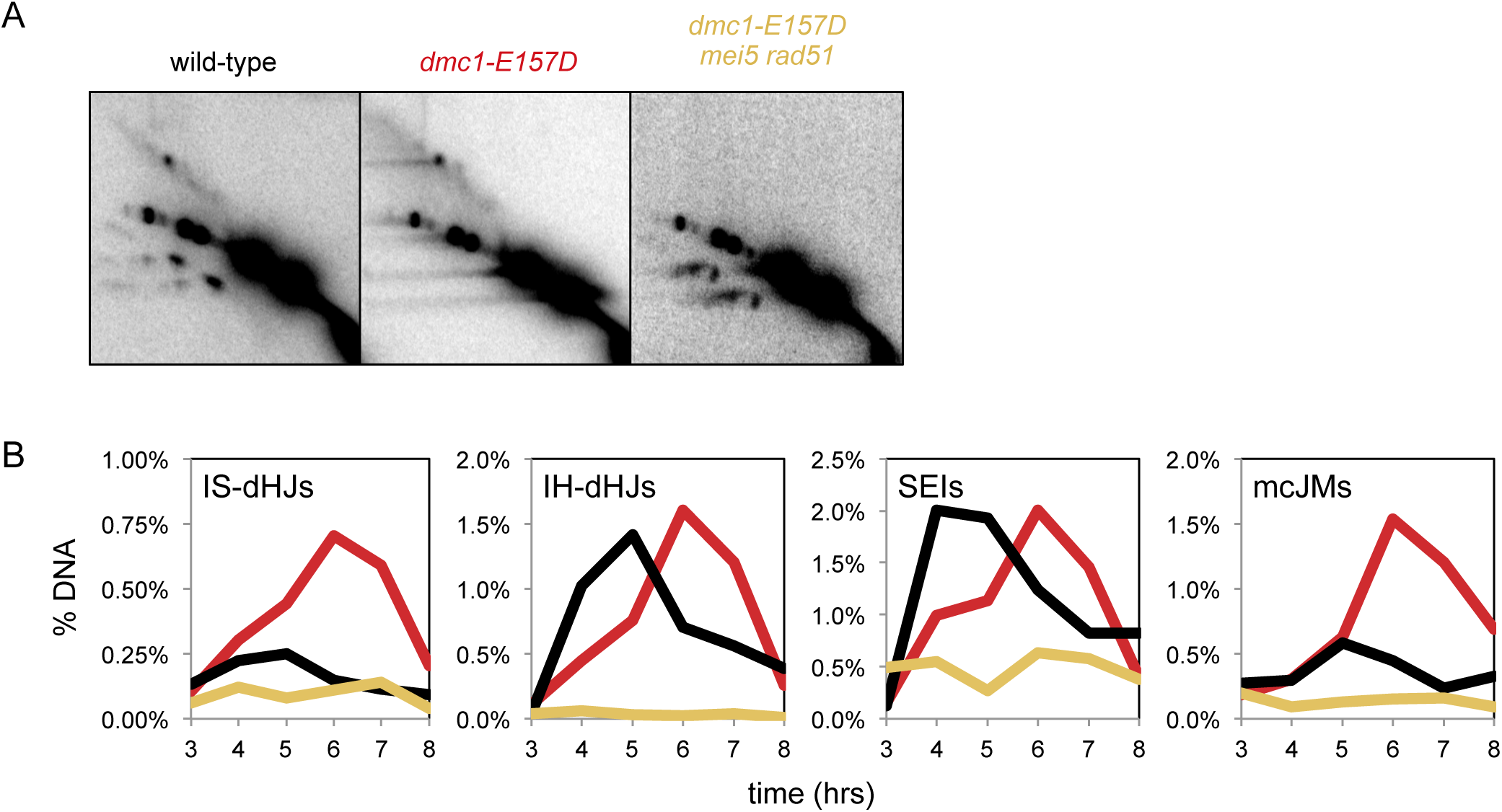
An independently derived diploid strain (DKB6413) corresponding to the *dmc1-E157D mei5 rad51* genotype gives the same result as shown in Figure 3. Wild-type and *dmc1-E157D* samples were prepared in parallel as controls. (a) 2D gels gels at the *HIS4::LEU2* hotspot from meiotic time course experiments. Representative images were chosen according to the time at which total JMs peaks for each sample. From left to right: wild-type (5h), *dmc1-E157D* (6h), *dmc1-E157D mei5 rad51* (7h). (b) 2D gel quantitation; black – wild-type, red – *dmc1-E157D*, yellow – *dmc1-E157D mei5 rad51.* Strains used in this experiment in the order in which they appear in figure, right to left: DKB3698, DKB6342, DKB6413.

**Supplemental Figure 4.**
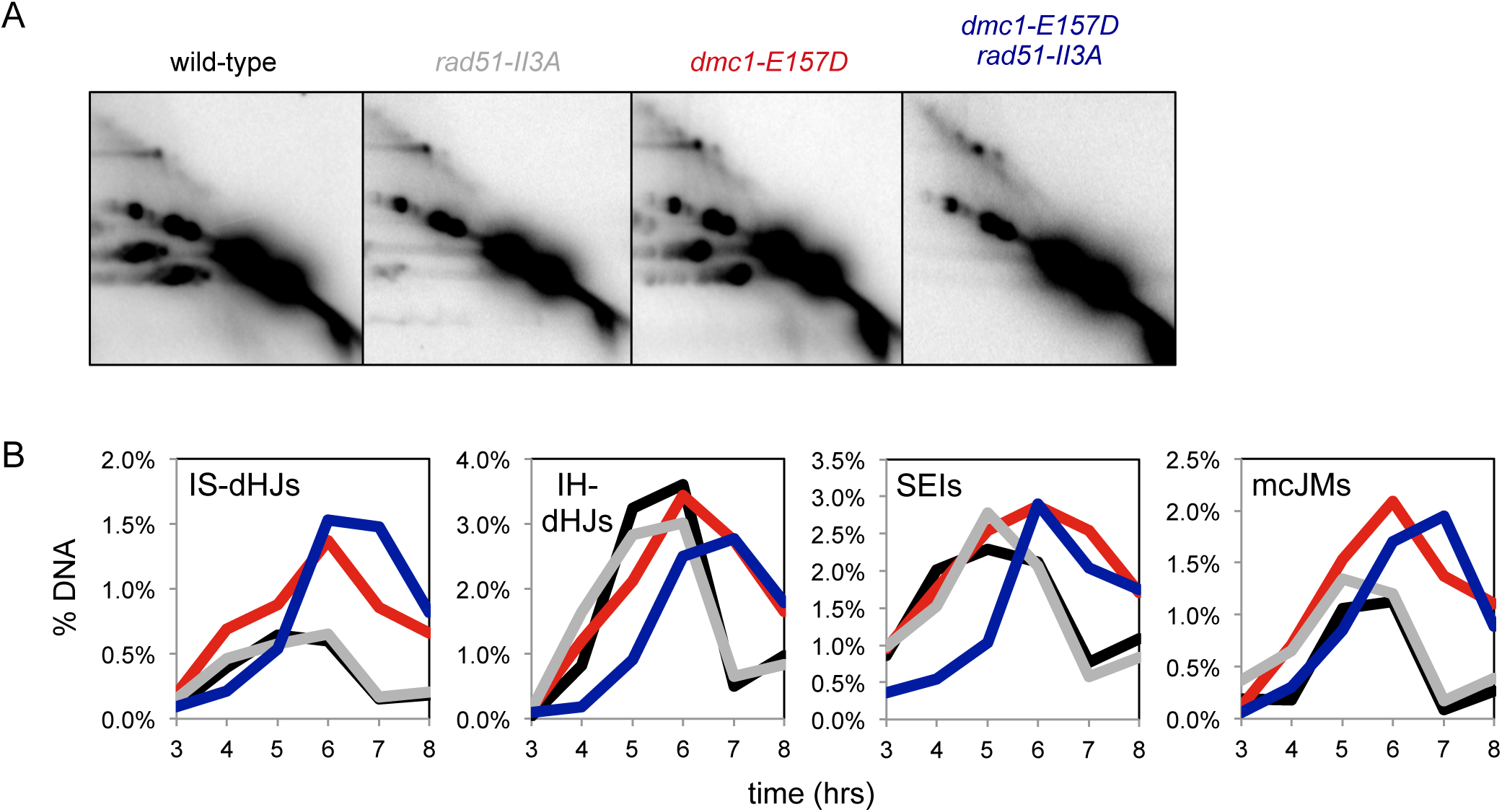
The defects associated with *dmc1-E157D rad51* are independent of Rad51’s catalytic activity. (a) 2D gels gels at the *HIS4::LEU2* hotspot from meiotic time course experiments. Representative images were chosen according to the time at which total JMs peaks for each sample. From left to right: wild-type (6h), *rad51-II3A* (6h), *dmc1-E157D* (6h), *dmc1-E157D rad51-II3A* (6h). (b) 2D gel quantitation; black – wild-type, gray – *rad51-II3A*, red – *dmc1-E157D*, dark blue – *dmc1-E157D rad51-II3A*. Strains used in this experiment in the order in which they appear in figure, right to left: DKB3698, DKB3689, DKB6342, DKB6400.

**Supplemental Figure 5.**
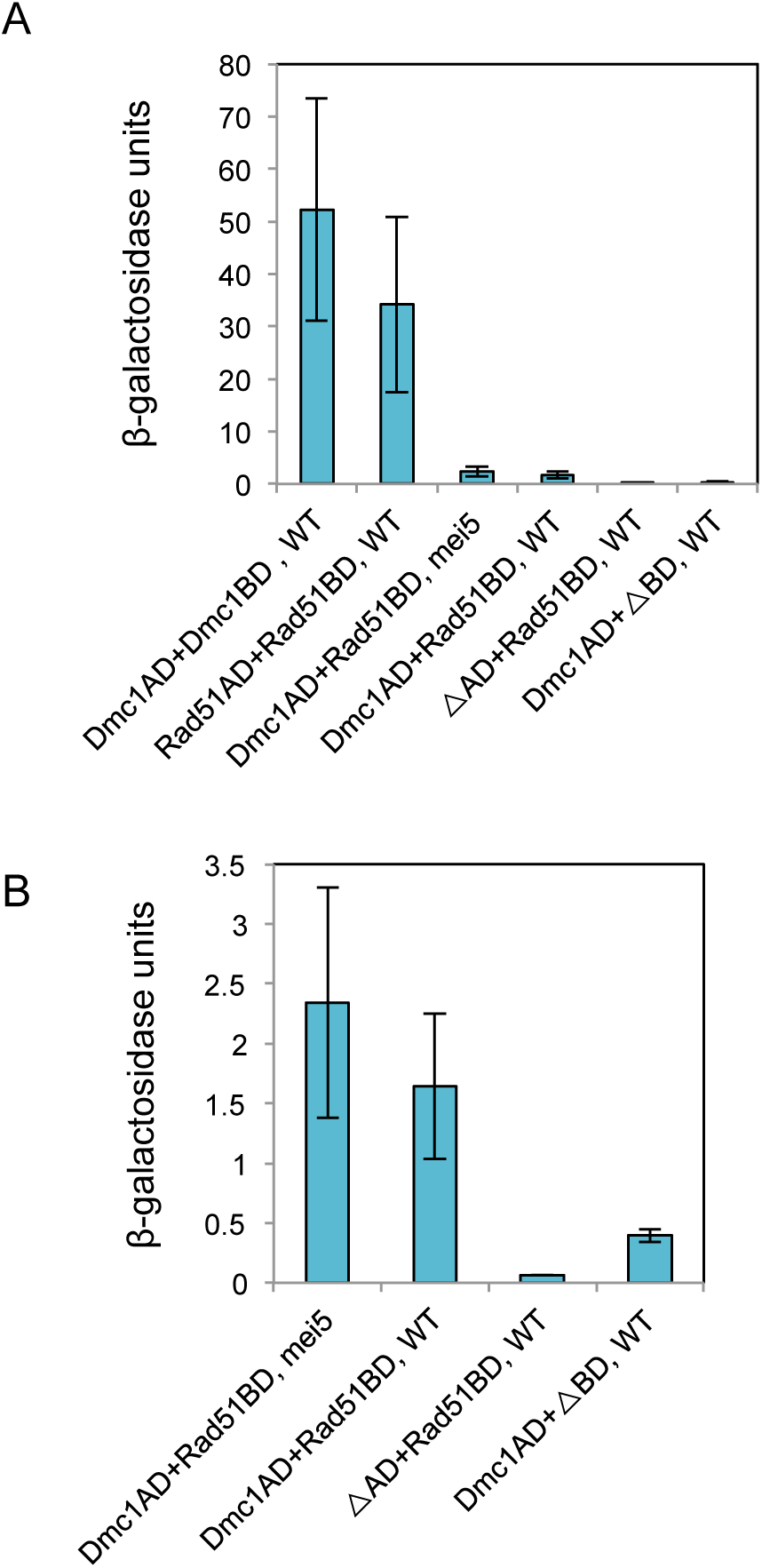
Meiotic two-hybrid analysis detects a weak interaction between Rad51 and Dmc1 that is independent of Mei5. (a) All interactions examined are plotted. (b) Subset of the same data shown in (a) to facilitate comparison of measurements of the Rad51-Dmc1 interaction with empty vector controls. ΔBD and ΔAD represent the empty vectors. Strains used in this experiment: DKB6501, DKB6503, DKB6508, DKB6509, DKB6513, DKB6515.

**Supplemental Figure 6.**
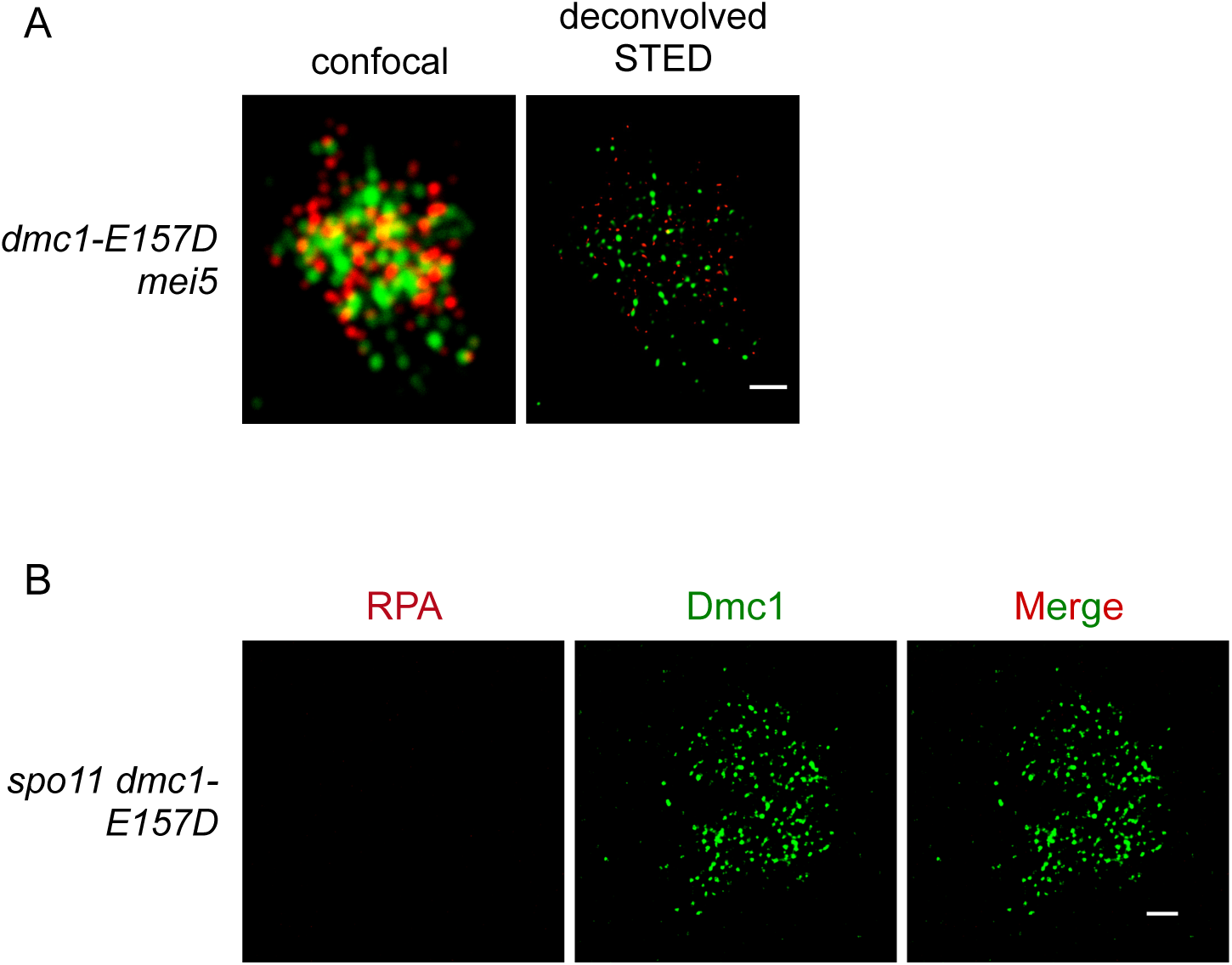
Super-resolution imaging resolves closely spaced foci, but elongated foci still form in *spo11 dmc1-E157D*. (a) Spread meiotic nuclei prepared from a *dmc1-E157D mei5* 5 hour sample imaged using confocal and STED microscopy methods. (b) STED imaging of a *spo11 dmc1-E157D* spread meiotic nuclei at 5 hours. For both, scale bar represents 1 micrometer. Red, RPA, green, Dmc1. Strains used in this experiment: DKB630, DKB6419.

**Supplemental Figure 7.**
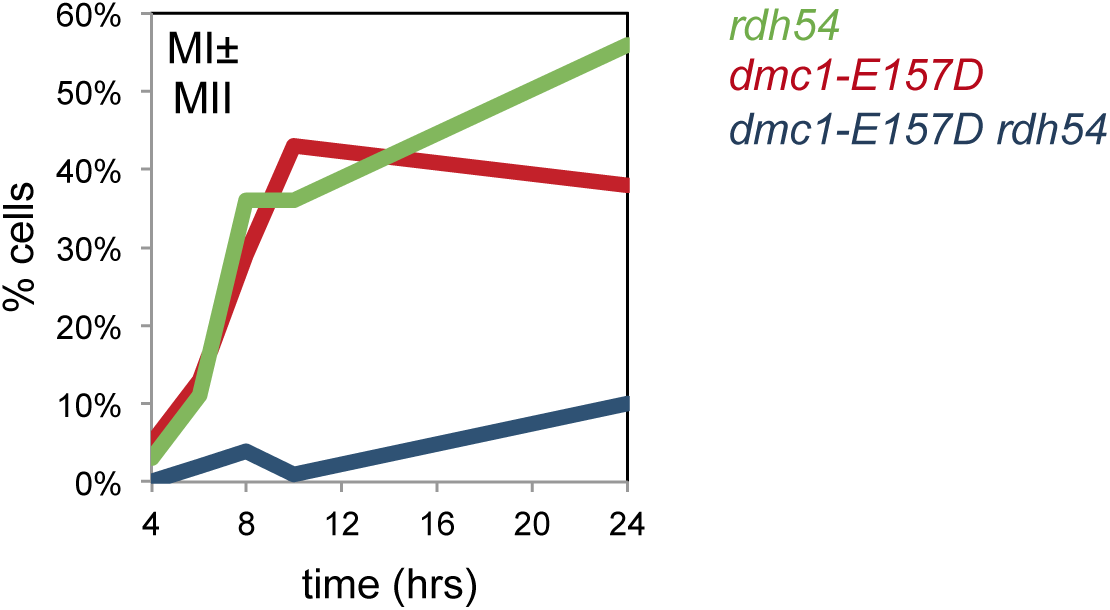
*dmc1-E157D rdh54* is more defective in meiotic progression than either of the single mutants, *dmc1-E157D* and *rdh54*. Meiotic progression data for strains indicated. For each time point, ≥100 cells were scored. Strains used in this experiment in the order in which they appear in figure, top to bottom: DKB2526, DKB6342, DKB6583.

**Supplemental Figure 8.**
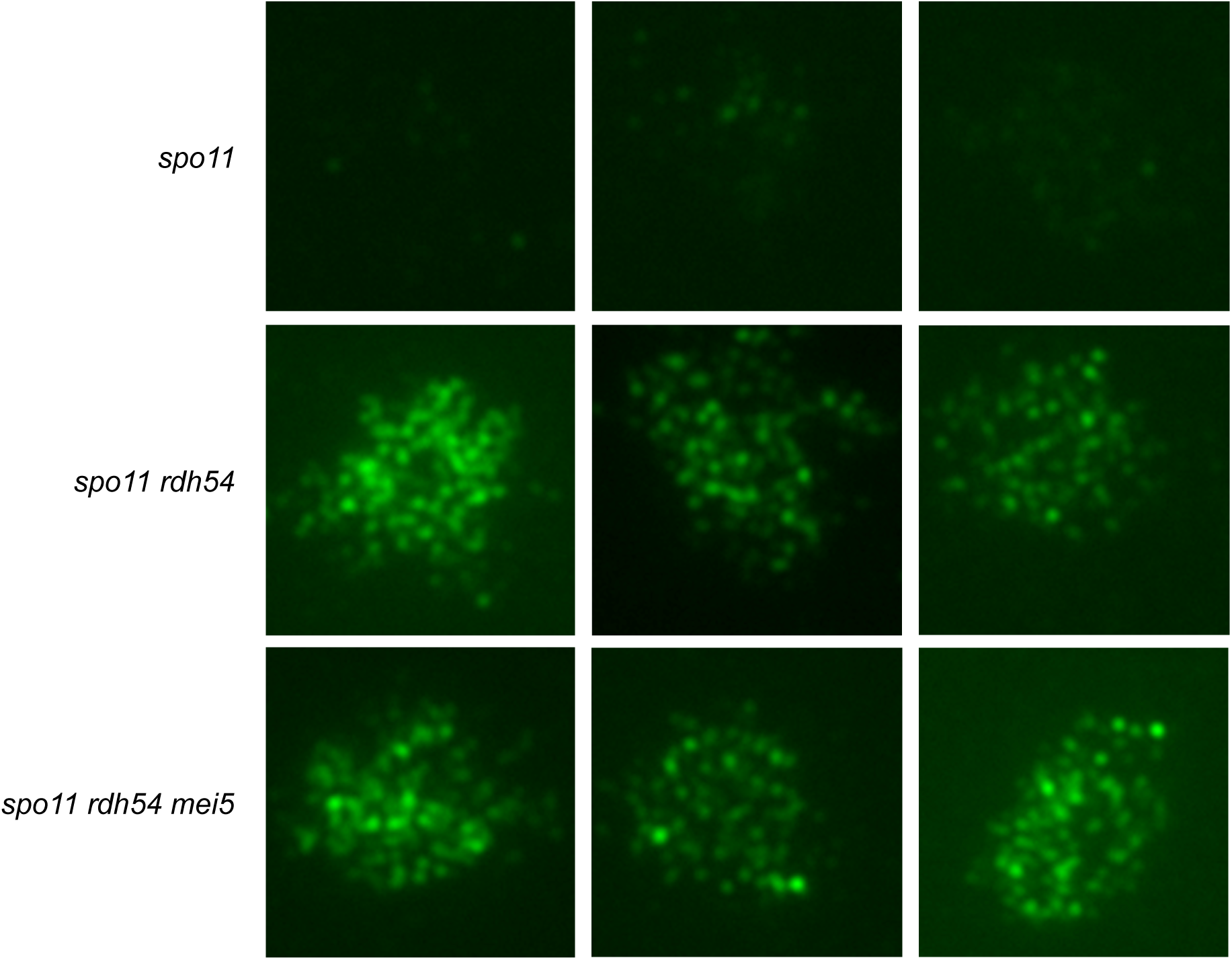
DSB-independent Dmc1-WT focus formation does not require Mei5. Samples were collected 4 hours after induction of meiosis in liquid medium and immuno-stained for Dmc1 and Hop2. Because Hop2 staining is Spo11 independent and specific for meiotic prophase, random prophase nuclei were selected on the basis of being Hop2 positive and then imaged for Dmc1 staining. 50 nuclei were examined for each sample with 3 representative nuclei shown for each of the three strains examined. Images were generated by wide-field microscopy using the same camera settings for all strains. Strains used in this experiment in the order in which they appear in figure, top to bottom: DKB2524, DKB2523, and DKB6571.

**Supplemental Table 1.**
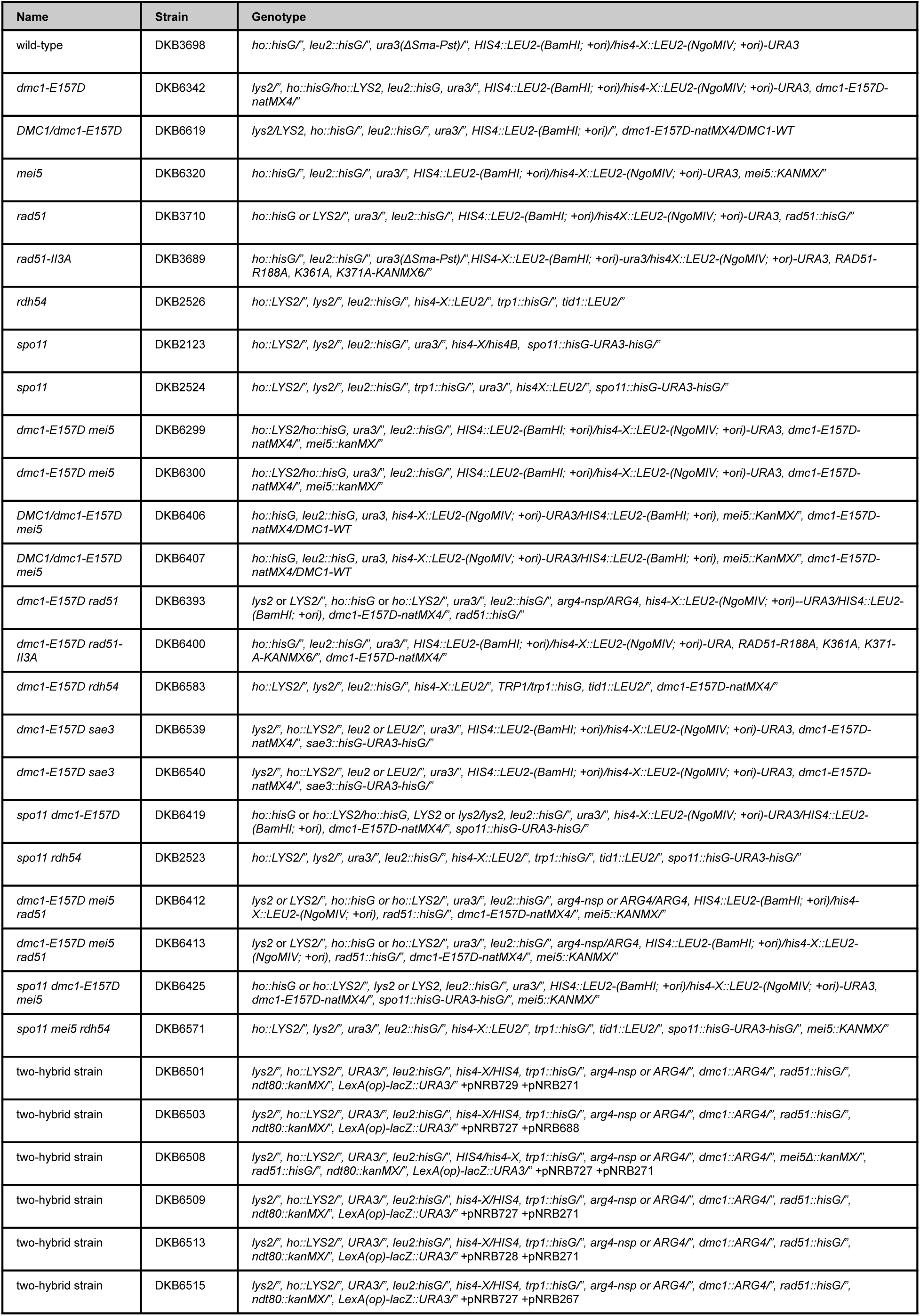
Yeast strains used in this study.

